# Interleukin-21 Drives a Hypermetabolic State and CD4^+^ T Cell-associated Pathogenicity in Chronic Intestinal Inflammation

**DOI:** 10.1101/2023.06.02.543518

**Authors:** Adebowale O. Bamidele, Shravan K. Mishra, Petra Hirsova, Patrick J. Fehrenbach, Lucia Valenzuela-Pérez, Hyun Se Kim Lee

**Affiliations:** Immunometabolism and Mucosal Immunity Laboratory, Division of Gastroenterology and Hepatology, Mayo Clinic, 200 First Street SW, Rochester, MN 55905, USA; Department of Immunology, Mayo Clinic, 200 First Street SW, Rochester, MN 55905, USA; Division of Gastroenterology and Hepatology, Mayo Clinic, 200 First Street SW, Rochester, MN 55905, USA

**Keywords:** Inflammatory Bowel Disease, Regulatory T cells, Mitochondria-ER appositions, Pyruvate, Interleukins

## Abstract

**BACKGROUND & AIMS:** Incapacitated regulatory T cells (Tregs) contribute to immune-mediated diseases. Inflammatory Tregs are evident during human inflammatory bowel disease (IBD); however, mechanisms driving the development of these cells and their function are not well understood. Therefore, we investigated the role of cellular metabolism in Tregs relevant to gut homeostasis.

**METHODS:** Using human Tregs, we performed mitochondrial ultrastructural studies via electron microscopy and confocal imaging, biochemical and protein analyses using proximity ligation assay, immunoblotting, mass cytometry and fluorescence-activated cell sorting, metabolomics, gene expression analysis, and real-time metabolic profiling utilizing Seahorse XF analyzer. We utilized Crohn’s disease single-cell RNA sequencing dataset to infer therapeutic relevance of targeting metabolic pathways in inflammatory Tregs. We examined the superior functionality of genetically-modified Tregs in CD4^+^ T cell-induced murine colitis models.

**RESULTS:** Mitochondria-endoplasmic reticulum (ER) appositions, known to mediate pyruvate entry into mitochondria via VDAC1, are abundant in Tregs. VDAC1 inhibition perturbed pyruvate metabolism, eliciting sensitization to other inflammatory signals reversible by membrane-permeable methyl pyruvate (MePyr) supplementation. Notably, IL-21 diminished mitochondria-ER appositions, resulting in enhanced enzymatic function of glycogen synthase kinase 3 β (GSK3β), a putative negative regulator of VDAC1, and a hypermetabolic state that amplified Treg inflammatory response. MePyr and GSK3β pharmacologic inhibitor (LY2090314) reversed IL-21-induced metabolic rewiring and inflammatory state. Moreover, IL-21-induced metabolic genes in Tregs *in vitro* were enriched in human Crohn’s disease intestinal Tregs. Adoptively transferred *Il21r*^-/-^ Tregs efficiently rescued murine colitis in contrast to wild-type Tregs.

**CONCLUSIONS:** IL-21 triggers metabolic dysfunction associated with Treg inflammatory response. Inhibiting IL-21-induced metabolism in Tregs may mitigate CD4^+^ T cell-driven chronic intestinal inflammation.

## Introduction

Persistent activation of diverse cytokine pathways in adaptive immune cells, such as CD4^+^ T cell subsets, is a hallmark of immune-mediated diseases, including human inflammatory bowel disease (IBD) ^1^. IBD, mainly Crohn’s disease (CD) and ulcerative colitis (UC), is a heterogenous and relapsing disorder characterized by gastrointestinal tissue damage and extraintestinal manifestations, with a high rate of surgical interventions ^2^. Despite therapeutic success of biologics against T cells, such as anti-tumor necrosis factor α (TNF-α), anti-p40 subunit shared by interleukin-12 (IL-12) and IL-23, and anti-integrin, inconsistent patient response remains a clinical concern ^3, 4^. Thus, our understanding of IBD is incomplete, highlighting the need to investigate critical drivers of inflammatory CD4^+^ T cells to uncover resistance mechanisms and maximize responsiveness.

Upon antigen-induced activation, naïve CD4^+^ T cells consume glucose, proliferate, and differentiate into functionally distinct regulatory and effector T helper (Th) cell lineages, as evidenced by unique transcription factor expression, metabolic programs and cytokine expression ^5^. In regulatory T cells (Tregs), transforming growth factor-beta 1 (TGF-β1) sustains the expression of Treg-defining forkhead domain-containing protein 3 (FOXP3) transcription factor necessary to subdue effector T cell-driven diseases ^6, 7^. Indeed, patients and mice lacking functional TGF-β1 develop multiorgan inflammation including IBD ^8, 9^.

Recent discovery of the interface between the immune system and metabolism is contributing immensely to our understanding of the complexity of immune cell regulation. Metabolic pathways differentially govern T cell differentiation and immune response, such as intracellular cytokine production and cell trafficking ^10^. For instance, activated CD4^+^ T cells favor glycolysis (conversion of glucose-derived pyruvate to lactate) and glutaminolysis, with decreased entry of pyruvate into mitochondria ^11, 12^. Pyruvate lies at the intersection between glycolysis and mitochondrial tricarboxylic acid (TCA) cycle, where immunometabolic hubs controlled by mitochondria-associated membranes (MAMs) are present ^13^. However, how micro-environmental cues control pyruvate metabolism in human Tregs is unclear. Metabolites not only promote the activity of the electron transport chain (ETC) but also enhance the generation of mitochondrial reactive oxygen species (mtROS) and mitochondrial DNA linked to signaling events and cellular function ^14, 15^. While several metabolic pathways, including glycolysis, underlie effector CD4^+^ T cell-mediated pathologies ^16^, it is unresolved whether metabolic dysfunction is associated with Treg abnormality reported during human IBD pathogenesis ^17–20^. Extracellular cues, such as TGF-β1, IL- 6, IL-12, IL-21 and IL-23, are known to control the balance between Treg and effector T cell response ^1, 21^. Yet how Tregs sense extracellular cues and integrate signals by fine-tuning MAMs to reshape extra-and intra-cellular metabolites, intracellular cytokines, cell migration and immune response in healthy and inflammatory conditions are poorly understood. Thus, we explored a possible role of MAMs, such as those enriched with the endoplasmic reticulum (ER), in responding to cues that control metabolic signaling events relevant to Treg immune-suppressive function.

Here, we showed that inhibition of pyruvate transport at mitochondria-associated ER membranes or the inner mitochondrial membrane (IMM) in induced Tregs (iTregs) triggered metabolic changes that resulted in sensitization to acquisition of an effector T cell-phenotype. Moreover, we identified IL-21 as the cytokine that uncoupled mitochondria from ER, resulting in enhanced glycogen synthase kinase 3 β (GSK3β) function, metabolic rewiring including glycolysis, pyruvate accumulation and exacerbation of IL- 12-induced inflammatory response. Like GSK3β pharmacologic inhibition, supplementation of iTregs with mitochondria membrane-permeable methyl pyruvate impaired glycolysis and inflammatory response triggered by IL-21 and IL-12. Overall, IL-21 exploited GSK3β, a metabolic checkpoint, to promote intracellular rewiring of iTregs. Notably, we found that IL-21-induced metabolic genes in iTregs were significantly enriched in intestinal Tregs of refractory IBD patients. In naïve CD4^+^ T cell-induced murine colitis, IL-21 receptor (IL-21R)-deficient Tregs were more efficient in alleviating disease severity than wild-type Tregs.

## Materials and Methods

Please see Supplementary Materials and Methods.

## Results

### Mitochondria-ER Appositions are Present in Human Tregs

Mitochondria and ER membranes can serve as immunometabolic hubs involved in facilitating pyruvate metabolism and mitochondrial respiration ^22, 23^. At mitochondria-ER junctions, voltage-dependent anion channel 1 (VDAC1) on the outer mitochondrial membrane (OMM) and inositol triphosphate receptor 1 (IP3R1) on the ER tethers mitochondria and ER membranes to facilitate mitochondrial influx of pyruvate ^13, 24, 25^. Yet limited data exist about how mitochondria-ER actively transform its ultrastructure to regulate Treg immune response. We therefore performed fundamental characterization of human Tregs to identify mitochondrial-ER appositions.

Induced Tregs (iTregs), well established to function *in vitro* and *in vivo* ^26^, were derived from naïve CD4^+^ T cells isolated from healthy human peripheral blood mononuclear cell (PBMC) donors (Figure 1A). Human iTregs and naïve control cells were subjected to transmission electron microscopy (TEM) to visualize mitochondria. Electron microscopy images showed that iTregs displayed a higher percentage of mitochondria in close association with ER than naïve cells where these interactions were minimal (59.04% vs. 3.27%, *P* < 0.0001, Figure 1B, C). Thus, electron-dense zones between the OMM and ER at contact sites in iTregs indicated that these were distinct subcellular structures (Figure 1B bottom panels – black arrows). TEM also revealed disparities in mitochondrial morphology. Human iTregs displayed long, tubular mitochondria associated with mitochondrial respiration ^27^, while mitochondria in naïve cells were rounded (mitochondrial aspect ratio per cell: 3.11 vs. 1.67, *P* < 0.0001 and aspect ratio per mitochondrion: 3.25 vs. 1.68, *P* < 0.0001, Supplementary Figure 1A-C). We substantiated our EM ultrastructural assessment with fluorescent-based organelle tracking in live cells. Imaging data showed higher mitochondria-ER co-localization in iTregs than naïve cells (Pearson’s coefficient of 0.58 vs. 0.39, *P* < 0.001, and 0.56 vs. 0.44, *P* < 0.01, using two different MitoTracker dyes, Supplementary Figure 1D, E). Fixed iTregs and CD4^+^ CD25^+^ CD127^dim/-^ T cells (human natural Tregs) were stained with VDAC1 and IP3R1 monoclonal antibodies, then subjected to proximity ligation assay (PLA) to biochemically quantify mitochondria-ER interactions. PLA can reliably detect protein-protein interactions of ≤ 30 nm distance *in situ* with high specificity and sensitivity. The discrete red PLA signals suggested that mitochondria-ER appositions were present in iTregs (Figure 1D, E) and natural Tregs (Supplementary Figure 1F).

**Figure 1.**
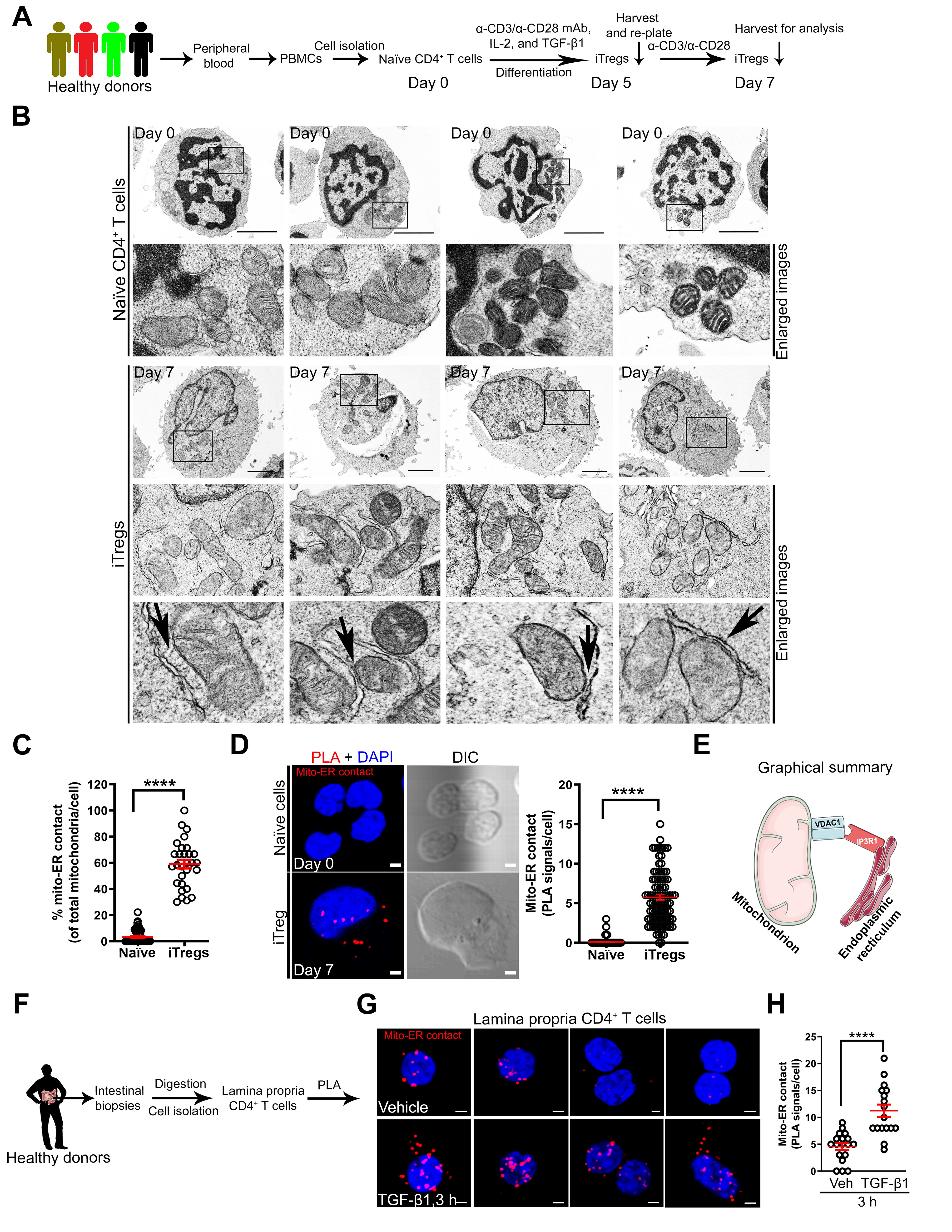
Mitochondria-ER Appositions are Present in Human Tregs. (A) Experimental workflow for naïve CD4^+^ T cell isolation from PBMCs and differentiation into iTregs. (B and C) Transmission electron microscopy (TEM) images of naïve cells (top panels) vs. iTregs (bottom panels) with enlarged images of mitochondria (mito)-ER contact. Black arrows indicate electron dense regions; scale bars, 2 µm (B). Graph shows mitochondria in contact with ER per cell in TEM images; naïve cells (n = 51) and iTregs (n = 29) (C). (D) Representative images show mito-ER contact in red after PLA, cell structure in differential interference contrast (DIC), and nucleus stained with 4’,6’-diamidino-2-phenylinode (DAPI) (DNA, blue); scale bar, 2 µm. Graph shows the number of PLA signals (mito-ER contact) per cell (n = 239 iTregs, n = 90 naïve cells). (E) Graphical summary of mitochondria-ER interaction detected via VDAC1-IP3R1 binding in human Tregs. (F) Experimental workflow for lamina propria (LP) CD4^+^ T cell isolation and PLA. (G) Representative images show mito-ER contact in red, DAPI in LP CD4^+^ T cells in the presence (bottom panels) or absence (top panels) of (±) TGF-β1 (25 ng/ml) stimulation; scale bar, 2 µm. (H) Graph shows quantitation of mito-ER contact (PLA signals) per LP CD4^+^ T cell; vehicle-stimulated cells (n = 17) and TGF-β1-stimulated cells (n = 18). Data represents mean ± SEM from 2-3 independent experiments or biological replicates. **** p < 0.0001, using two-tailed Student’s t test.

Healthy lamina propria (LP) CD4^+^ T cells known to exhibit Treg-like phenotype and respond to TGF-β1 ^28, 29^ were isolated from human intestinal biopsies and examined for mitochondria-ER interaction via PLA (Figure 1F). We found mitochondria-ER interactions in a subset of LP CD4^+^ T cells (Figure 1G top panels), and these interactions were enhanced by TGF-β1 (4 vs. 11 PLA signals per cell, *P* < 0.0001, Figure 1G, H). IP3R1 interaction was specific to VDAC1 in contrast to translocase of the outer mitochondrial membrane 20 (TOM20) where TOM20-IP3R1 binding was minimal (98% vs. 5% PLA^+^ LP CD4^+^ T cells, *P* < 0.0001, Supplementary Figure 1G), which supports the specificity of this interaction. Using complementary approaches in human samples, mitochondria-ER interactions – detected via VDAC1-IP3R1 binding – are present in human Tregs.

### VDAC1 Inhibition Alters iTreg Metabolic State and Sensitizes to IL-12-induced Inflammatory Response

Previous study has demonstrated the capacity to generate functional iTregs in the absence of TGF-β1^30^. We showed VDAC1-IP3R1 interactions, i.e., mitochondria-ER appositions, were more abundant in iTregs generated in the presence of TGF-β1 than iTregs derived in the absence of TGF-β1 on day 5 (*P* < 0.001) and 7 (*P* < 0.001) as shown Figure 2A-C. As controls, we showed successful development of FOXP3^+^ cells in both cell types (Supplementary Figure 2A), while TGF-β1-induced CD103 (αE) integrin ^31^ confirmed cellular responsiveness to TGF-β1 (Supplementary Figure 2B). Mitochondria-ER interaction has been linked to mitochondrial pyruvate transport and oxidative phosphorylation (OXPHOS) ^13, 22^. Consistent with these reports, oxygen consumption rate (OCR) was higher in iTregs generated in the presence of TGF-β1) than iTregs derived in the absence of TGF-β1, as evidenced by increased basal, ATP-coupled and FCCP-induced maximal (Figure 2D). In summary, abundant mitochondria-ER interactions correlate with higher OCR, which we found to be less enriched in effector Th1, Th2, and Th17 cells (Figure 2E and Supplementary Figure 2C). We then explored whether perturbing mitochondria-ER function, i.e., pyruvate transport, would impact OCR and Treg immune response.

**Figure 2.**
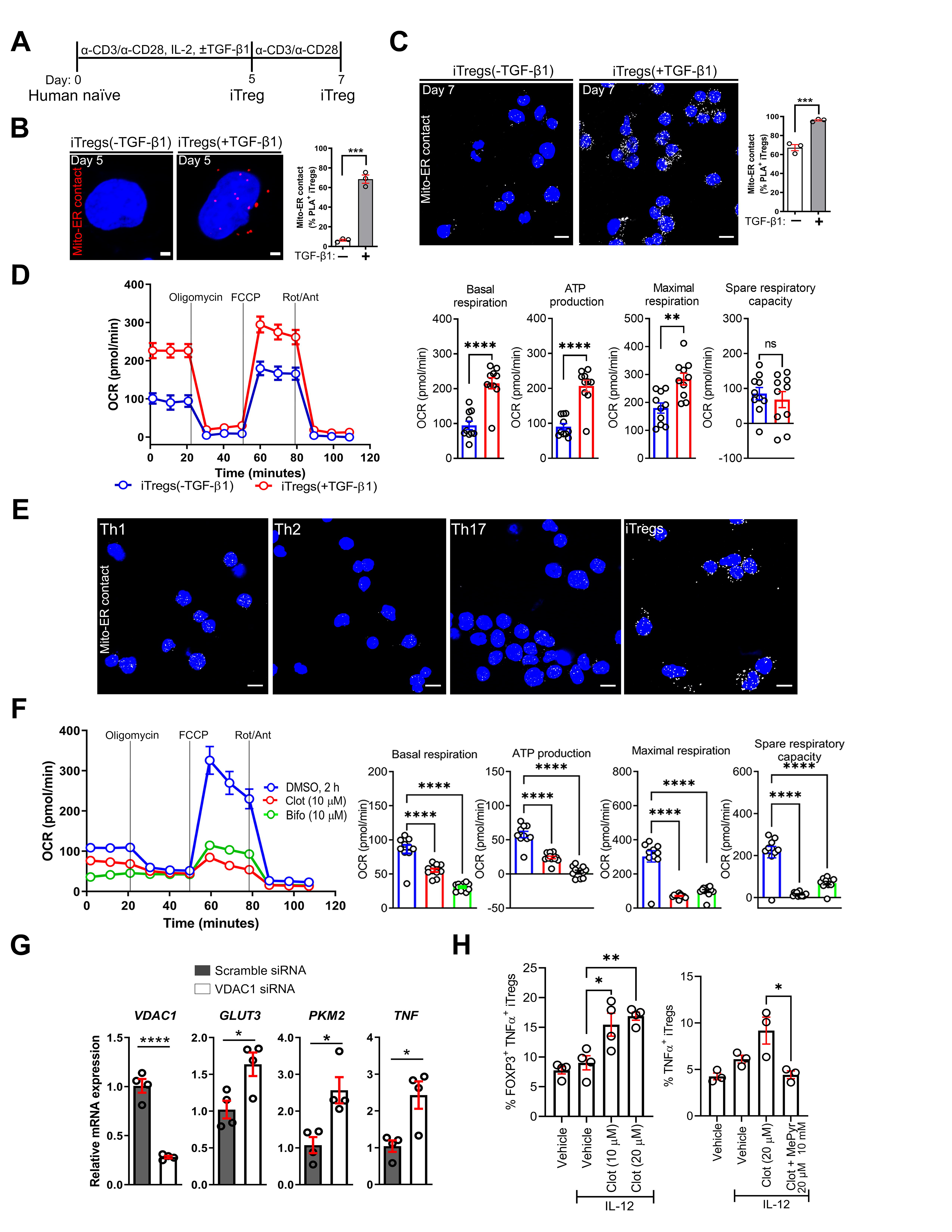
VDAC1 Inhibition Alters iTreg Metabolic State and Sensitizes to IL-12-induced Inflammatory Response. (A) Experimental workflow for the generation of human iTregs in the absence or presence of TGF-β1. (B and C) Representative PLA images of iTregs ± TGF-β1 show mito-ER contacts in red on day 5 (B) or in white on day 7 (C); scale bars, 2 µm and 10 µm. Graphs show PLA^+^ cells on day 5 (n = 257 vs. n = 200) (B) and on day 7 (n = 129 vs. n = 119) (C). (D) Representative oxygen consumption rate (OCR) profile of iTreg cell types before and after mitochondrial perturbation (n = 3). Bar graphs show calculated basal respiration, ATP production, maximal respiration, and spare respiratory capacity; mean ± SEM from 10-12 technical replicates. (E) Representative PLA images show mito-ER contact in iTregs vs. effector T helper cells (n = 3 biological donors); scale bar, 10 µm. (F) Representative OCR profile of iTregs ± clotrimazole (clot), bifonazole (bifo) or UK5099 (n = 3). Bar graphs show calculated basal respiration, ATP production, maximal respiration, and spare respiratory capacity; mean ± SEM from 12 technical replicates. (G) Relative mRNA expression in transfected iTregs via RT-qPCR (n = 3). (h) Percentage of TNF-α^+^ in iTregs, as determined by flow cytometry (n = 3-4). Data represents mean ± SEM. * p < 0.05, ** p < 0.01, *** p < 0.001, and **** p < 0.0001, using two-tailed Student’s t test or one-way ANOVA followed by Bonferroni test for multiple comparisons.

At the OMM, mitochondria-ER interactions stabilize hexokinase (HK)-I localization, thereby activating VDAC1 and directing pyruvate into the IMM via the mitochondrial pyruvate carrier (MPC) ^13, 32, 33^. Thus, we reasoned that VDAC1 inhibition in iTregs would diminish pyruvate transport and OCR. Consistent with our hypothesis, acute inhibition (2 h) of HK-I-mediated activation of VDAC1 with clotrimazole or bifonazole ^34^, herein referred to as clot and bifo (10-20 µM), reduced OCR (basal OCR, ATP-linked OCR, FCCP-induced maximal OCR, space respiratory capacity: *P* < 0.0001, Figure 2F). Similarly, UK5099 (2-20 µM, 2 h)-mediated destabilization of MPC protein complexes decreased OCR (ATP-linked OCR: *P* < 0.05, FCCP-induced maximal OCR: *P* < 0.0001, spare respiratory capacity: *P* < 0.0001) Supplementary Figure 2D). Of note, siRNA-mediated depletion of VDAC1 increased the mRNA expression of *TNF* (*P* < 0.05) and glycolysis genes, such as glucose transporter 3 (*GLUT3*; *SLC2A3*, *P* < 0.0001) and pyruvate kinase M 2 (*PKM2, P* < 0.05) (Figure 2G). In agreement with VDAC1 depletion, clotrimazole treatment sensitized cells to IL-12, a known activator of signal transducer and activator of transcription 4 (STAT4) and Th1 signature TNF-α and IFN-γ cytokines, as evidenced by increased percentage of TNF-α^+^ iTregs (Figure 2H left). Mechanistically tying this inflammatory response to impaired mitochondrial pyruvate transport, supplementation with methyl pyruvate (MePyr, 10 mM), a mitochondria membrane-permeable metabolite (Figure 2H right), reversed clotrimazole-mediated TNF-α production. Like VDAC1 inhibitor, UK5099 increased the percentage of IL-12-induced IFN-γ^+^ iTregs, with IFN-γ^+^ iTregs enriched in T cell homing CD49D (α4) integrin (Supplementary Figure 2E). By utilizing various approaches, we inferred that interfering with pyruvate transport at the OMM or IMM induced metabolic rewiring that sensitized iTregs to acquiring an effector T cell-like phenotype (Supplementary Figure 2F). These experiments mechanistically linked pyruvate transport to metabolic homeostasis that limits iTreg inflammatory response. We then explored whether extracellular cues could trigger metabolic alterations and inflammatory responses by altering mitochondria-ER ultrastructure.

### IL-21 Stimulation of Human iTregs Promotes a Hypermetabolic State

IL-21 can enhance IFN-γ and IL-17 expression in LP T cells from human IBD subjects ^35, 36^, mirroring MPC inhibition. We then hypothesized that exposure of iTregs to IL-21 would induce metabolic alterations linked to inflammatory response. Indeed, Central Metabolism PCR Array and RT-qPCR demonstrated that IL-21 independently upregulated transcripts associated with glycolysis and effector T cell response (Figure 3 and Supplementary Figure 3). These include *SLC2A3* (*GLUT3*), hexokinase 2 (*HK2*), glucose phosphate isomerase (*GPI*), enolase 1/2 (*ENO1/2*), glyceraldehyde 3-phosphate dehydrogenase (*GAPDH*), *PKM2*, monocarboxylate transporter 4 (*MCT4,* also known as *SLC16A3*), and hypoxia-inducible factor 1 alpha (*HIF1A*) (Figure 3A-C, 3F). IL-21 also amplified the citrate-to-acetyl-CoA shuttle system previously associated with glycolytic-lipogenic metabolism in Th17 cells ^37^. These include citrate synthase (*CS*), citrate transporter (*SLC25A1*), ATP citrate lyase (*ACYL*), acetyl-CoA carboxylase 1 (*ACC1; ACACA*), and fatty acid synthase (*FASN*) (Figure 3A, D, E). IL-21 transcriptionally activated fatty acid oxidation (FAO) and glutaminolysis known to support the TCA cycle, OXPHOS and anabolic demands (Figure 3A, B), including glutamate oxaloacetate transaminase 1 (*GOT1*) previously linked to 2-hydroxyglutarate (2-HG) generation and Th17 cell commitment ^38^. IL-21 did not have a significant effect on *FOXP3* and *TGFB1* mRNA (Figure 3F, 3G). In summary, profiling of IL-21-stimulated iTregs suggests a metabolic rewiring to glycolysis and other compensatory pathways (Figure 3H). IL-21-induced metabolism was associated with inflammatory response, as evidenced by elevated *IFNG*, *IL17A* and *IL17F* (*P* < 0.05, Figure 3I) while there was no reduction the percentage of cells expressing FOXP3 and other markers (Figure 3J).

**Figure 3.**
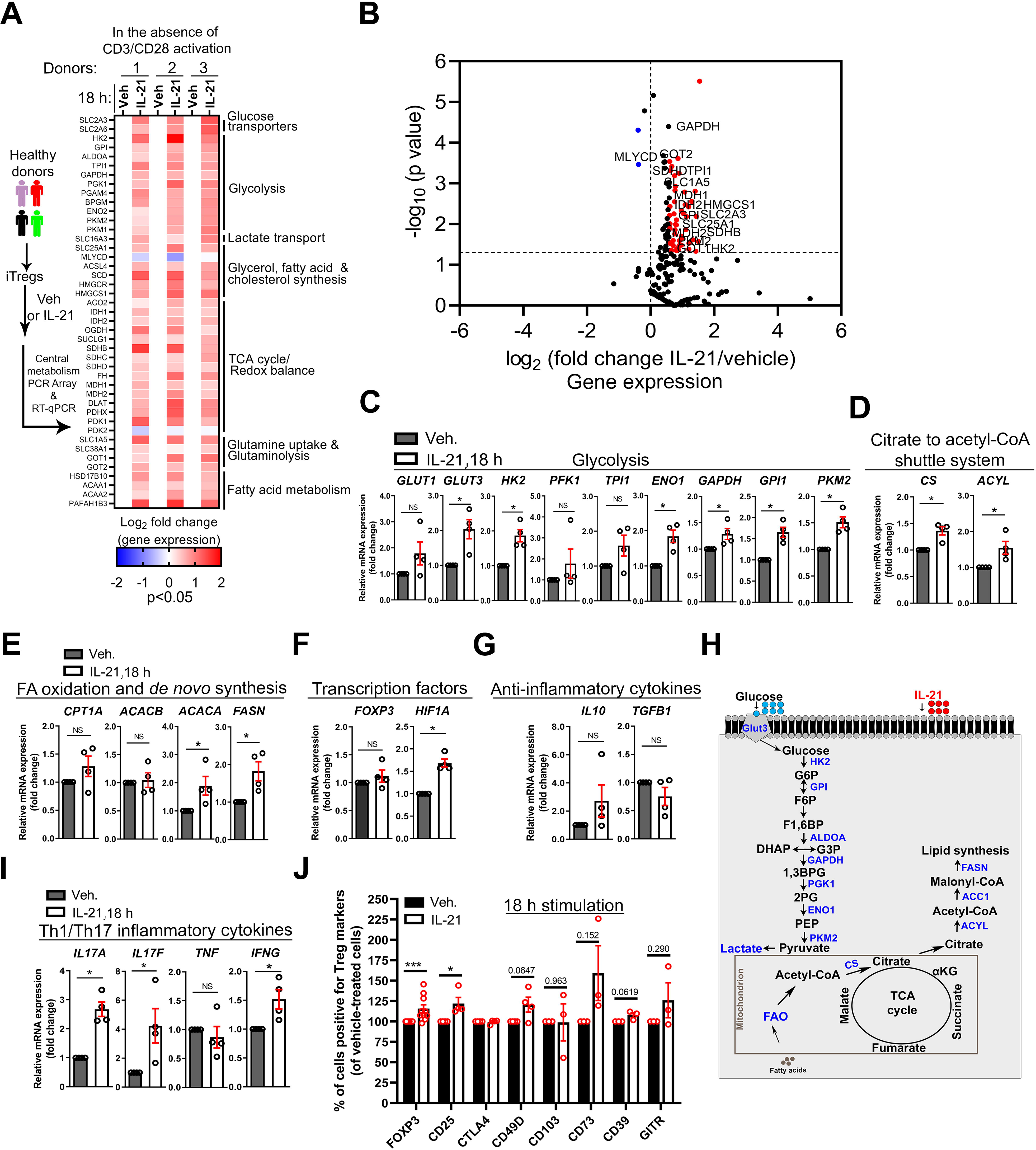
IL-21 Stimulation of Human iTregs Promotes a Hypermetabolic State. (A-I) Transcriptional profiling of human iTregs ± IL-21 (100 ng/ml) in serum-free media in the absence of TCR and CD28 activation. (A) *Left*, experimental workflow for the metabolic transcriptional profiling of iTregs. *Right*, heatmap of 43 metabolic transcripts (IL-21/vehicle, p < 0.05) and the associated metabolic processes (n = 3). (B) Volcano plot illustrates all detected 211 metabolic genes (IL-21/vehicle, n = 3). Red dots denote genes with >1.5-fold change and p-value of <0.05 while blue dots denote <-1.0-fold change and p-value of <0.05. (C-G) Relative mRNA expression via RT-qPCR in iTregs ± IL-21 (n = 4). Expression of metabolic genes (C-E), transcription factors (F), and anti-inflammatory cytokines (G); Mann-Whitney *U* test. (H) Schematic illustrates metabolic genes upregulated in IL-21-stimulated iTregs. (I) Relative mRNA of inflammatory genes in iTregs via RT-qPCR ± IL-21 (n = 4); Mann-Whitney *U* test. (J) Percentage of iTregs positive for FOXP3 and surface markers, as determined by FACS (n = 3-9). Data represents mean ± SEM. * p < 0.05 and *** p < 0.001, using two-tailed Student’s t test. NS (not significant).

### IL-21 Dissociates Mitochondria from ER, Resulting in Pyruvate Imbalance and Sensitization to IL- 12-induced Inflammatory Response

Addressing a mechanism by which IL-21 induced these transcriptional changes, we found that Tregs exposed to IL-21 (2 h) displayed reduced mitochondria-ER appositions, with altered mitochondrial morphology and volume, as determined by PLA, TEM, live-cell imaging, and serial block-face scanning EM (SEM) (Figure 4A-C and Supplementary Figure 4A, B). Glucose-regulated protein 75 (GRP75) tethers VDAC1 to IP3R1^13, 39^, and in agreement, IL-21 reduced GRP75 co-localization with mitochondria (Supplementary Figure 4C). Unlike IL-21, other control cytokine IL-6, IL-23 and IL-12 failed to disrupt mitochondria-ER interaction within the time point evaluated (Figure 4A, Supplementary Figure 4D), indicating IL-21 as a potential negative regulator of mitochondria-ER coupling. Mitochondria-ER interaction is functionally coupled to GSK3β inactivation (via phosphorylation), and subsequent VDAC1 activation and pyruvate entry into mitochondria ^13, 33, 40^. We found that IL-21 reduced phosphorylated GSK3β (pSer9-GSK3β), the inactive form (*P* < 0.01 by immunoblot, *P* < 0.05 by FACS, Figure 4D and Supplementary Figure 4E). We and others interpret this loss of inactive form as a gain in GSK3β enzymatic activity, given the unspecific nature of antibodies raised against GSK3β active moiety. Increased pTyr705-STAT3 indicated cellular responsiveness to cytokines (Figure 4D). We then speculated that IL-21-induced mitochondria-ER defect would decrease intracellular ATP, due to impaired mitochondrial pyruvate metabolism, and indeed, ATP was expectedly reduced (*P* < 0.05, Figure 4E). Thus, our data suggest that IL-21 uncouples mitochondria from ER, resulting in GSK3β activation and disruption of ATP production.

**Figure 4.**
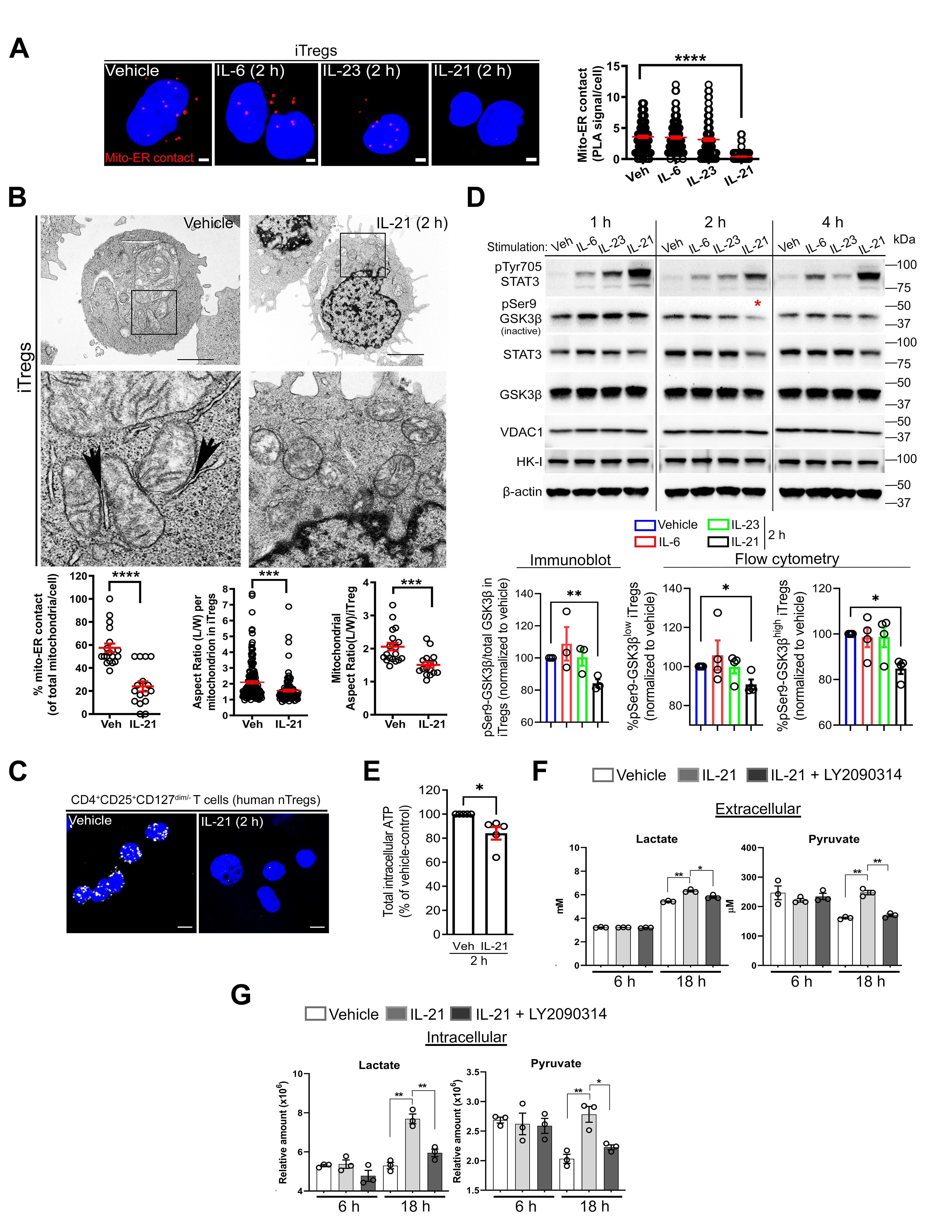
IL-21 Dissociates Mitochondria from ER, Resulting in Pyruvate Imbalance and Sensitization to IL-12-induced Inflammatory Response. (A) Representative PLA images of mito-ER contact in iTregs ± 50 ng/ml of IL-6 (n = 113 cells), IL-23 (n = 131 cells), or IL-21 (n = 221 cells) vs. vehicle (n = 115 cells); scale bar, 2 µm. Graph shows mito-ER contact (red PLA signals) per cell (n = 2 independent experiments). (B) Representative TEM images of iTregs ± IL-21, with black arrows indicating electron dense regions; scale bar, 2 µm. Graphs show quantitation of mito-ER contact (n = 19 vs. n = 16 cells) (left), aspect ratio per mitochondrion (n = 167 vs. n = 101 mitochondria) (middle), and mitochondrial aspect ratio per cell (n = 19 vs. n = 16 cells) (right). (C) Representative PLA images show mito-ER contact in human nTregs treated with vehicle or IL-21 after activation and TGF-β1 stimulation for 24 h; scale bar, 5 µm (n = 2 independent experiments). (D) Representative immunoblots from iTregs ± cytokines (100 ng/ml). Red asterisk shows reduction in GSK3β inactive form (top) (n = 3). Bar graphs show pSer9-GSK3β levels by immunoblot (n = 3) and the percentage of pSer9-GSK3β-expressing cells by FACS (n = 4). (E) Intracellular ATP levels in iTregs ± IL-21 (50 ng/ml) (n = 5); Mann-Whitney *U* test. (F and G) Extra-and intra-cellular lactate and pyruvate abundance from iTregs cultured in normal conditions ± IL-21 (100 ng/ml), LY2090314 GSK3 inhibitor (0.025 µM) or both (n = 3). Data represents mean ± SEM. * p < 0.05, ** p < 0.01, *** p < 0.001 and **** p < 0.0001, using two-tailed Student’s t test or one-way ANOVA followed by Bonferroni test for multiple comparisons.

To understand how GSK3β enzymatic activity is linked to IL-21-induced nutrient metabolism, we quantified the abundance of metabolites under standard culturing conditions. Extracellular pyruvate and lactate were more abundant in the media collected from IL-21-stimulated iTregs (6-18 h) than in vehicle-treated cells (Figure 4F); however, excreted glutamine-derived glutamate was less abundant (6 h), with mininal changes in glucose levels (Supplementary Figure 4F). IL-21 increased intracellular lactate and pyruvate levels (Figure 4G) as well as glycolysis intermediates and TCA cycle metabolites (Supplementary Figure 4G), which is consistent with data in Figure 3A. In summary, IL-21-induced mitochondria-ER defect is associated with dysregulation of extra-and intra-cellular lactate and pyruvate.

To mechanistically link GSK3β activity and pyruvate-lactate imbalance to inflammatory cytokine expression, we hypothesized that pharmacologic inhibition of active GSK3β would restore metabolite levels and consequently suppress inflammatory cytokine expression. Consistent with our hypothesis, treatment with pharmacologic inhibitor of GSK3α and GSK3β isoforms (LY2090314) ^41^, herein referred to as LY (0.025-0.1 µM), reversed IL-21-induced metabolite imbalance, including lactate and pyruvate levels (Figure 4F, G, and Supplementary Figure 4F, G). We then examined the anti-inflammatory effect of LY2090314 in the same experimental context as our VDAC1 and MPC inhibition studies by utilizing iTregs exposed to IL-21 and IL-12 as our model system, given that IL-21 also sensitized cells to IL-12- induced IFN-γ production (Supplementary Figure 4H). LY2090314 and the inhibitor of the rate-limiting step in glycolysis (2-deoxy-Ɒ-glucose, 2-DG, 10 µM, Supplementary Figure 4I) reduced the percentage of IFN-γ^+^ and TNF-α^+^ iTregs (Supplementary Figure 4J), which is consistent with Figure 3A and 4F, G. Using established pharmacological inhibitors ^16^, we examined the contribution of other impacted metabolic pathways to inflammatory cytokine production (Supplementary Figure 4I). ETC complex I and III inhibitors (rotenone and antimycin A, respectively) and ATP uncoupler (FCCP) diminished the percentage of IFN-γ^+^ and or TNF-α^+^ iTregs by various extent (Supplementary Figure 4J), which is consistent with the emerging role of mitochondria in mediating T cell effector response^42^. On the contrary, PKM2 activator (TEPP-46) and FAO, ACYL, ACC1, TCA cycle enzyme and ETC complex II and V inhibitors had no significant effect on the percentage of IFN-γ^+^ and TNF-α^+^ iTregs (Supplementary Figure 4J, K). Collectively, IL-21 disrupts mitochondria-ER interaction and potentiates GSK3β function, resulting in metabolic alterations that promote IL-12-induced IFN-γ and TNF-α.

### Methyl Pyruvate Supplementation Mirrors LY2090314 Treatment and Suppresses the Metabolic Basis of Inflammatory iTregs

We then comprehensively measured changes in OCR and ECAR (extracellular acidification rate due to lactate production, i.e., rate of glycolysis) to mechanistically investigate metabolic adaptations occurring in real time in response to IL-21 stimulation (1-48 h). As verified in glucose concentrations (1-10 mM), IL- 21 (1 h) reduced OCR (Figure 5A, B), coinciding with mitochondria-ER decoupling and GSK3β activation (Figure 3A-D). Consistent with data in Figure 3A, carnitine palmitoyltransferase 1A (CPT1A)-mediated initiation of long-chain FAO potentially contributed to IL-21-induced OCR (3-18 h) (Figure 5C, D), as CPT1A inhibitor, etomoxir (ETO, 50 µM, 3 h), restored OCR (Figure 5D). Thus, IL-21 induces OCR fluctuations, mirroring Th17 cells in which OXPHOS supports IL-17 production ^43–45^.

**Figure 5.**
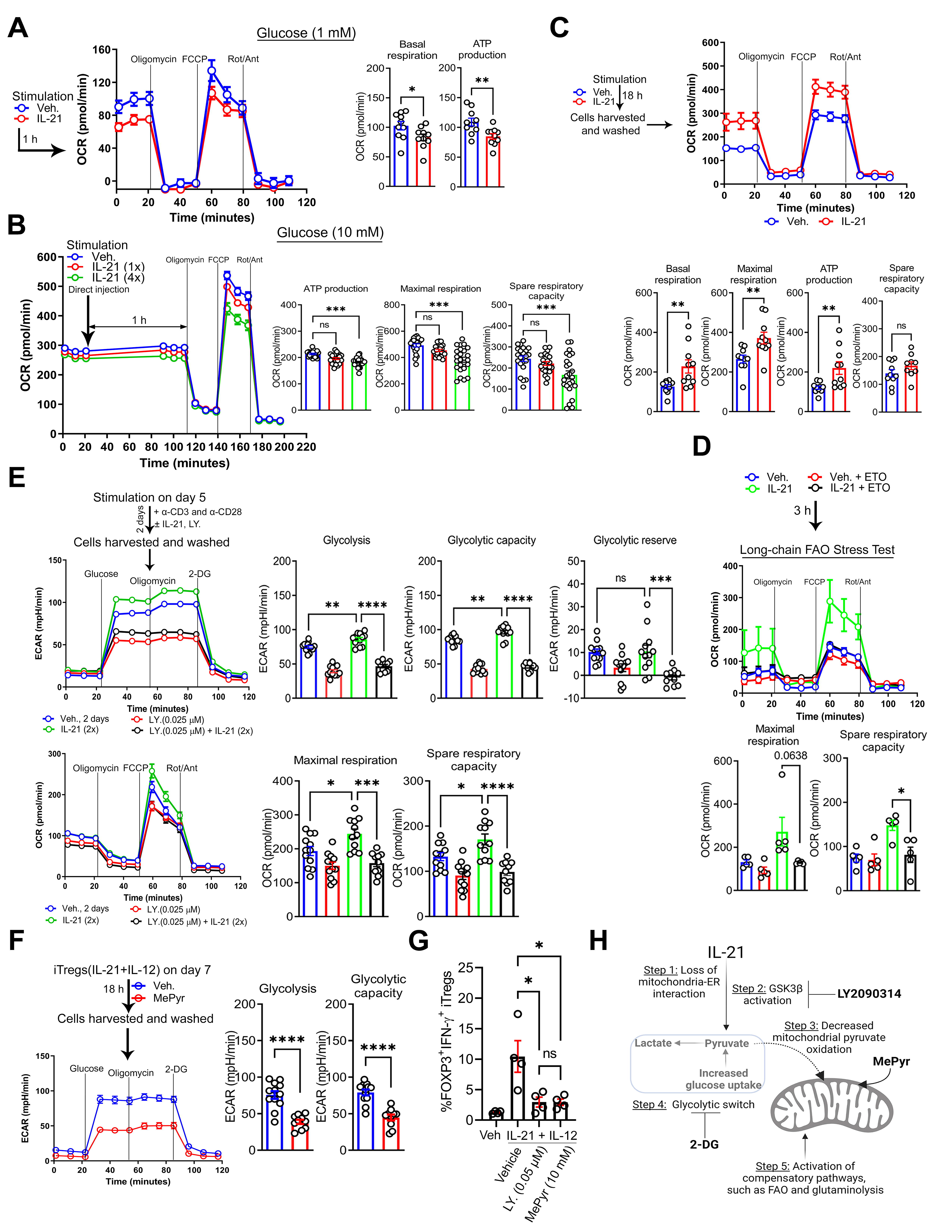
Methyl Pyruvate Supplementation Mirrors LY2090314 Treatment and Suppresses the Metabolic Basis of Inflammatory iTregs. (A) Representative OCR profile of iTregs in 1 mM glucose ± IL-21 (100 ng/ml) (n = 3). Bar graphs show calculated basal respiration and ATP production; mean ± SEM from 10 technical replicates. (B) Representative OCR profile of iTregs in 10 mM glucose ± IL-21 (100-400 ng/ml) (n = 3). Bar graphs show calculated basal respiration, ATP production, maximal respiration, and spare respiratory capacity; mean ± SEM from 10-24 technical replicates. (C) Representative OCR profile of iTregs ± IL-21 (100 ng/ml) in the presence of TCR and CD28 activation (n = 3). Bar graphs show calculated basal respiration, maximal respiration, ATP production and spare respiratory capacity; mean ± SEM from 10 technical replicates. (D) Representative OCR profile of iTregs ± IL-21 (100 ng/ml), etomoxir (ETO, 50 µM) or both (n = 3). Bar graphs show calculated basal respiration, maximal respiration, ATP production and spare respiratory capacity; mean ± SEM from 5 technical replicates. (E) Representative ECAR (top) and OCR (bottom) profiles of iTregs ± IL-21, LY2090314, or both (n = 3). Bar graphs show calculated glycolysis, glycolytic capacity, and glycolytic reserve (top) and maximal respiration and spare respiratory capacity (bottom); mean ± SEM from 10-12 technical replicates. (F) Representative ECAR profile of IL-21 and IL-12-stimulated iTregs ± methyl pyruvate (MePyr, 10 mM) (n = 3). Graphs show calculated glycolysis and glycolytic capacity; mean ± SEM from 10-12 technical replicates. (G) Graph shows percentage of IL-21 and IL-12-induced FOXP3^+^ IFN-γ^+^ iTregs ± LY2090314 or MePyr (n = 4). (H) Schematic diagram illustrates IL-21-mediated metabolite imbalance in iTregs. Diagram was created with biorender.com Data represents mean ± SEM. * p < 0.05., *** p < 0.001 and **** p < 0.0001, using two-tailed Student’s t test or one-way ANOVA followed by Bonferroni test for multiple comparisons.

Notably, LY2090314 reversed ECAR and OCR induced by IL-21 (Figure 5E). Consistent with data in Figure 4J, mass cytometry by Time of Flight (CyTOF) showed that LY2090314 reduced IL-21 and IL-12-induced IFN-γ, TNF-α and IL-17A across CD49D subpopulations (Supplementary Figure 5A, B). LY2090314 also reversed ECAR and or OCR induced by IL-6, IL-23, and UK5099 (Supplementary Figure 5C, D). These notable findings suggest that several proinflammatory cytokines can induce glycolysis, potentially via distinct mechanisms. Furthermore, GSK3β, by shuttling between its active and inactive forms, may act as a central metabolic checkpoint, which in turn dictates pyruvate localization (cytosolic vs. mitochondria) as supported by evidence from other cell types ^13, 33^.

Collectively, these mechanistic observations led us to then hypothesize that circumventing mitochondria-ER defect or requirement by merely supplementing IL-21 and IL-12-stimulated Tregs with membrane-permeable MePyr would reduce ECAR and inflammatory response, mirroring GSK3β inhibition. Indeed, MePyr diminished ECAR of IL-21 and IL-12-stimulated iTregs (basal glycolysis, glycolytic capacity: *P* < 0.0001, Figure 5F), and it reduced the percentage of IFN-γ^+^ iTregs, mirroring LY2090314 treatment (*P* < 0.05, Figure 5G). Overall, our work suggests that enforced mitochondrial pyruvate entry via MePyr supplementation or LY2093014 treatment suppresses the metabolic phenotype of inflammatory iTregs (Figure 5H), resulting in reduced inflammatory response. Next, we examined the immune-suppressive capacity of IL-21R-deficient (*Il21r*^-/-^) Tregs in alleviating murine colitis in comparison to wild-type (WT) Tregs.

### IL-21-induced Metabolic Genes are Enriched in Refractory Human IBD, and IL-21R-deficient Tregs Effectively Lessen CD4^+^ T Cell-induced Colitis in Mice

We first examined whether IL-21-induced metabolic genes enriched in our *in vitro* iTregs (Figure 3A) were represented in Tregs from severe lesions of human IBD. We analyzed single-cell RNA-sequencing (scRNA-seq) dataset from human CD ileal lesions (Martin et. al), in which a pathogenic cell-cell landscape consisting of Ig<u>G</u> plasma cells, inflammatory <u>m</u>ononuclear phagocytes, <u>a</u>ctivated <u>T</u> cells, and <u>s</u>tromal cells (termed GIMATS^high^ module) predicts patient resistance to anti-TNF therapy ^20^. Notably, in Treg clusters from inflamed tissues of GIMATS^high^ patients, we found higher expression of genes associated with glycolysis (*GAPDH*, *ENO1*, *ALDOA*, *LDHB*, and *TPI1*), nucleotide metabolism (*CMPK1*, *GUK1*, and *APRT*), FAO (*PAFAH1B3*), and glutaminolysis (glutamine transporter *SLC38A1*) than that of GIMATS^low^ patients (Figure 6A). Furthermore, some of these metabolic genes were also upregulated in Treg clusters from inflamed IBD tissues than that of adjacent uninflamed tissues from the same patients (Figure 6B). In summary, IL-21-induced metabolic genes in iTregs *in vitro* are enriched in intestinal Tregs derived from refractory IBD patients.

**Figure 6.**
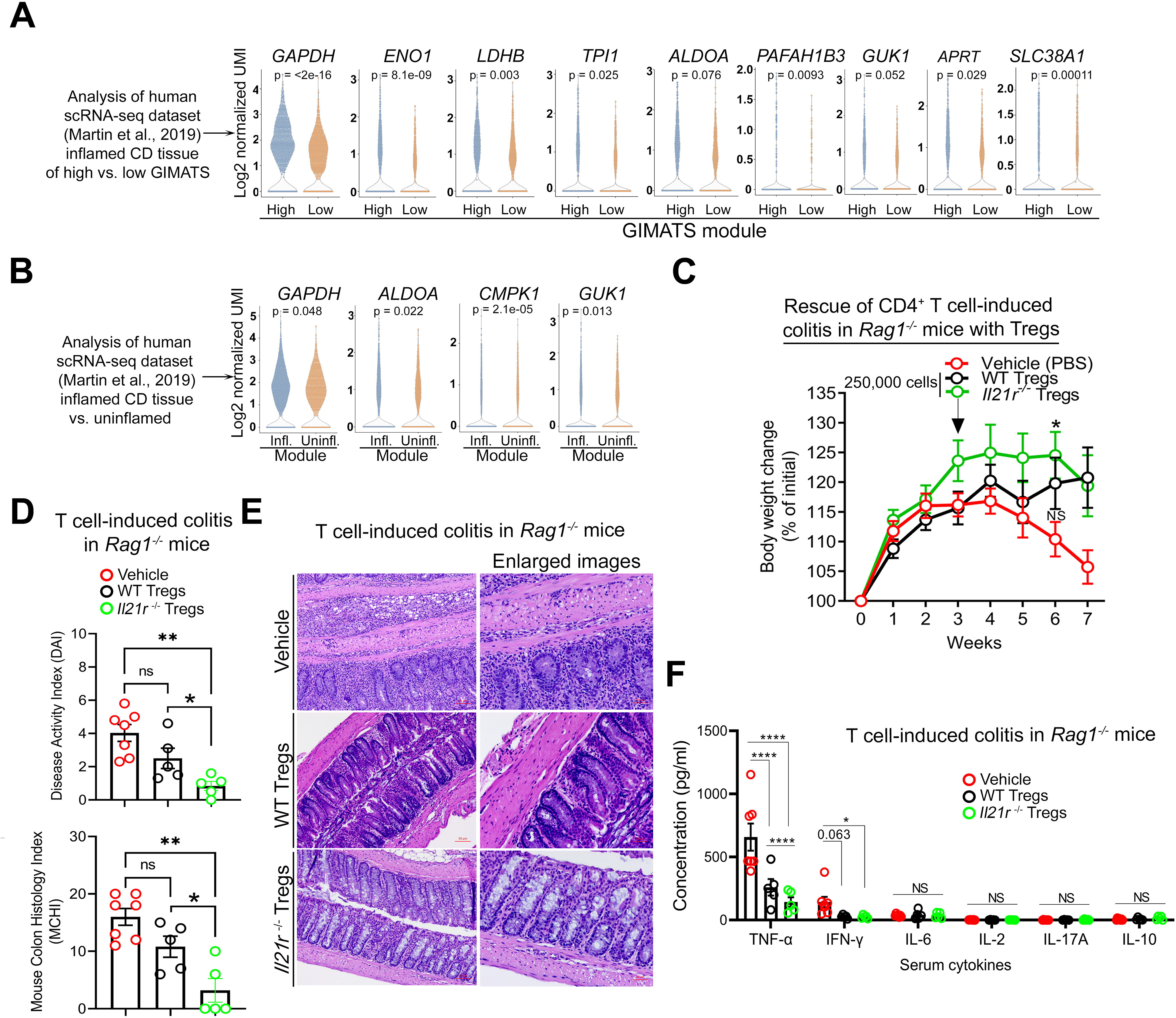
IL-21-induced Metabolic Genes are Enriched in Refractory Human IBD, and IL-21R-deficient Tregs Effectively Lessen CD4^+^ T Cell-induced Colitis in Mice. (A) Violin plots show the log2 normalized UMI of metabolic genes in human Treg clusters derived from Crohn’s disease (CD) inflamed tissues of GIMATS (Ig<u>G</u> plasma cells, inflammatory <u>M</u>NP, and <u>a</u>ctivated <u>T</u> and <u>s</u>tromal cells) module (Wilcoxon test; *P*); n = 5 for GIMATS^high^ patients and n = 4 for GIMATS^low^ patients. (B) Violin plots show the log2 normalized UMI of metabolic genes in human Treg clusters derived from CD inflamed vs. adjacent non-inflamed tissues from same patients (n = 9 patients) (Wilcoxon test; *P*). (C) Change in body weight of mice from week 1-7 during naïve CD4^+^ T cell-induced colitis progression in mice (n = 5-7 mice per group). (D) DAI (top), and MCHI (bottom) of treated colitis mice on week 7. MCHI was assessed by a blinded pathologist. (E) Serum cytokine expression analysis of treated colitis mice on week 7. (F) Hematoxylin and eosin (H&E) staining of colon sections of treated colitis mice on week 7; scale bar, 50 µm. Data represents mean ± SEM. * p < 0.05, ** p < 0.01, and **** p < 0.0001, using two-way ANOVA followed by Tukey for multiple comparisons (C), non-parametric Kruskal-Wallis test followed by Dunn’s multiple comparisons (vehicle vs. Tregs) and two-tailed Student’s t test (WT vs. *il21r*^-/-^ Tregs) (D)

Next, we hypothesized that *Il21r*^-/-^ Tregs would resolve colitis more efficiently than WT Tregs. We adoptively transferred naïve CD4^+^ CD45RB^high^ T cells into recombinase activating gene-1 (*Rag1*)^-/-^ mice to induce colitis, followed by injection with either vehicle, WT Tregs or *Il21r*^-/-^ Tregs on day 0 to prevent disease (Supplementary Figure 6A) or on week 3 to rescue disease (Figure 6C). Colitis mice injected with vehicle expectedly lost 15-20% of their initial weight (Figure 6C and Supplementary Figure 6A), displayed increased disease activity index (DAI), and developed colitis based on blinded histology assessment (mouse colitis histology index (MCHI) scoring) (Figure 6D and Supplementary Figure 6B). In the colitis rescue approach, *Il21r*^-/-^ Tregs successfully stabilized mice weight (*P* < 0.05, Figure 6C), reduced DAI (*P* < 0.05, Figure 6D top), contained established tissue inflammation and damage (Figure 6D bottom, 6E) and reduced serum TNF-α (*P* < 0.0001) in contrast to WT Tregs (Figure 6F right). In the colitis prevention approach, although both WT and *Il21r*^-/-^ Tregs improved weight gain (Supplementary Figure 6A) and reduced DAI (Supplementary Figure 6B left), WT Tregs failed to efficiently restrain tissue inflammation and damage (*P* > 0.05, Supplementary Figure 6B right and 6C) as well as serum IL-6 (*P* > 0.05, Supplementary Figure 6D). Taken together, using these complementary approaches, IL-21R-deficient Tregs exhibit enhanced capacity in suppressing the levels of inflammatory mediators and in resolving murine colitis induced by pathogenic CD4^+^ T cells.

## Discussion

Although intestinal Tregs expressing inflammatory cytokines in CD and UC patients have been described ^18–20^, the mechanism responsible for this inflammatory phenotype and its contribution to disease are poorly understood. Our prominent findings include the identification of mitochondria-ER appositions in human Tregs, which we believe facilitate pyruvate transport and its metabolism in the mitochondria. Impairing mitochondrial pyruvate entry by targeting mitochondria-ER architecture induces metabolic dysfunction and renders iTregs susceptible to acquiring an effector T cell-like phenotype. IL-21 uniquely perturbs mitochondria-ER ultrastructure and exploits GSK3β to promote Treg metabolism that favors inflammatory immune response. GSK3β inhibition or MePyr supplementation potentially bypasses mitochondria-ER requirement to suppress iTreg inflammatory response and its metabolic basis. Perturbations to other mitochondria-associated membranes could therefore be a generalizable mechanism pertinent to gut pathophysiology and other chronic inflammatory conditions. Our study indicates that GSK3β is perhaps constitutively active – acting as a central metabolic checkpoint at the pyruvate bifurcation point – given that its pharmacologic inhibition affected basal TNF-α, ECAR and OCR in normal iTregs. Adoptive transfer of IL-21R-deficient Tregs more efficiently ameliorates disease onset and progression, which mechanistically supports the relevance of IL-21-IL-21R axis *in vivo*. Overall, our study suggests that the ability of pathogenic cues to instigate metabolic alterations that result in Treg dysfunction might induce resistance to current monotherapies.

We showed that TGF-β1 and IL-21 had opposing effects on mitochondria-ER interaction. We suspect this may involve re-organization of actin and microtubule networks, with subsequent changes to the subcellular localization of chaperones such as GRP75. IL-21 stimulation activated several metabolic pathways, including pathways supportive of OXPHOS. Although mitochondrial metabolism is important for Treg function under steady-state conditions ^46^, mtROS and consequent DNA damage have been linked to Treg cell death and autoimmune pathogenesis ^47^. However, in our study, IL-21-induced mitochondrial alterations were associated with IFN-γ and TNF-α production. Presumably, disease-specific microenvironmental factors control metabolic rewiring and Treg cell fate decisions (inflammatory vs. cell death). Along this line, mitochondrial abnormality in intestinal epithelial cells has been associated with IBD ^48, 49^. Our study raises the possibility that IL-21 may broadly instigate mitochondrial dysfunction in multiple cell types, leading to metabolic alterations culminating in chronic inflammation.

We found that the prototypical glycolysis inhibitor (2-DG) significantly decreased IL-21 and IL-12- induced IFN-γ and TNF-α production by iTregs; however, the precise glycolytic mechanisms are not understood. It might involve proteins, such as HIF1α, GAPDH, mechanistic target of rapamycin, GLUTs and MCTs previously linked to effector T cell function ^5^. Furthermore, the contribution of metabolic pathways, such as one carbon and nucleotide metabolism, to Treg biology warrants to be investigated. Paradoxically, we found an upward trend in IL-10 expression in iTregs in response to IL-21 and IL-12, perhaps due to enhanced cholesterol biosynthesis previously linked to IL-10 induction ^50^. Of note, our inflammatory iTreg *in vitro* model mirrors data from inflamed tissues of UC patients in which FOXP3^+^ IL10^+^ T cells were also the primary source of TNF-α ^19^. We speculate that IL-21-induced IL-10 is an attempt to dampen IL-17 expression, given that IL-10 signaling can suppress IL-17 expression ^51^. Thus, our inflammatory iTregs *in vitro* resemble FOXP3^+^ T cells reported in IBD patients ^19, 20^. Elucidating how IL-21-induced inflammatory cytokines act in an autocrine manner to further compromise Treg function might reveal mechanisms pertinent to IBD. IL-21 reduced CD49D (α4), nevertheless, CD49D^high^ iTregs maintained TNF-α expression, consistent with the notion that highly inflammatory T cells possess gut-infiltrating capability. IL-21 may regulate Treg trafficking, with implications in intestinal tissue damage and extraintestinal manifestations; however, experiments are needed to substantiate this speculation. Interestingly, a higher percentage of α4β7^low^ Tregs was observed in CD patients ^17^, and the restoration of α4β7 in expanded *ex vivo* Tregs is being explored for autologous transplant to treat CD patients (NCT03185000). This implies that human Tregs, supported by evidence in murine models, also require α4β7 for selective homing to the gut for immunosuppressive function.

Overall, our studies uncovered a link between IL-21 and mitochondria-ER dysfunction, which results in enhanced GSK3β function, hypermetabolic state, and iTreg inflammatory response. We also provided several therapeutic strategies for halting Treg inflammatory state. Repurposing GSK3 inhibitor (LY2090314) for pharmacological intervention or generation of autologous Tregs might be beneficial. Remarkably, LY2090314 was well tolerated in a Phase 2 study for cancer treatment ^41^. Our findings provide the basis for future characterization of inflammatory Tregs that may inspire the development of immunomodulatory therapies.

## Abbreviations

FOXP3: forkhead domain–containing protein 3
IMM: inner mitochondrial membrane
OMM: outer mitochondrial membrane
MAM: mitochondria-associated membranes
VDAC1: voltage-dependent anion channel 1
IP3R1: inositol triphosphate receptor 1
MPC: mitochondrial pyruvate carrier
Mito-ER: mitochondria-endoplasmic reticulum
EM: electron microscopy
MePyr: methyl pyruvate
GSK3β: glycogen synthase kinase 3 beta
IBD: inflammatory bowel disease
CD: Crohn’s disease
UC: ulcerative colitis
IL: interleukin
IP: intraperitoneally
iTreg: induced regulatory T cell
mRNA: messenger RNA
OXPHOS, FCCP: carbonyl cyanide-p-trifluoromethoxyphenylhydrazone; oxidative phosphorylation
FAO: fatty acid oxidation
TCA: tricarboxylic acid
ECAR: extracellular acidification rate
OCR: oxygen consumption rate
ETC: electron transport chain
PBMC: peripheral blood mononuclear cell
LP: lamina propria
PLA: proximity ligation assay
PBS: phosphate-buffered saline
RNA-seq: RNA sequencing
TCR: T cell receptor
CD3: cluster of differentiation 3
CD28: cluster of differentiation 28
CD49D: integrin alpha 4
GLUT3: glucose transporter 3
PKM2: pyruvate kinase M2
TGF-β1: transforming growth factor beta 1
Th: T helper
TNF-α: tumor necrosis factor
IFN-γ: interferon gamma
iTreg: induced regulatory T cell
nTreg: natural regulatory T cell
WT: wild-type.

## Acknowledgements

We thank Mayo Clinic Microscopy and Cell Analysis Core and Mayo Clinic Immune Monitoring Core for their experimental and technical assistance.

## Transcript Profiling

GEO database under accession code GSE224033 (https://www.ncbi.nlm.nih.gov/geo/query/acc.cgi?acc=GSE224033)

## Author contributions

A. O. B., conceptualization, investigation, supervision, project administration, writing – original draft & funding acquisition; S. K. M., investigation; P. H., investigation, writing – review & editing, & resources; P. J. F., L. V., H. K., investigation.

