## Supplementary methods for "Interleukin-21 Drives a Hypermetabolic State and CD4^+^ T Cell-associated Pathogenicity in Chronic Intestinal Inflammation"

**Short title**: IL-21 induces metabolic disturbance in Tregs

Adebowale O. Bamidele[1, 2], Shravan K. Mishra[1], Petra Hirsova[3], Patrick J. Fehrenbach[1], Lucia Valenzuela-Pérez[3], Hyun Se Kim Lee[3]

[1]Immunometabolism and Mucosal Immunity Laboratory, Division of Gastroenterology and Hepatology, Mayo Clinic, 200 First Street SW, Rochester, MN 55905, USA
[2]Department of Immunology, Mayo Clinic, 200 First Street SW, Rochester, MN 55905, USA
[3]Division of Gastroenterology and Hepatology,
Mayo Clinic, 200 First Street SW, Rochester, MN 55905, USA

### Materials and Methods

*Healthy Blood Donors*

After written informed consent, we obtained blood samples from healthy male donors (19-60 years old) as buffy coats from apheresis cones (Mayo Clinic Blood Donor Center).

*Human intestinal biopsy specimens*

We obtained de-identified biopsy specimens from healthy donors. Before inclusion in this study, we received written informed consent from participants.

*Animals*

C57BL/6J *Rag1*$^{-/-}$ mice were purchased from the Jackson Laboratory and kept in conventional housing in the Mayo Clinic animal facility. C57BL/6NJ WT and *Il21r*$^{-/-}$ (*Il21r*$^{tm1kopf}$) [22] mice were purchased from the Jackson Laboratory. All mice were housed in the Mayo Clinic animal facility. Mice used in experiments were males of 6–9 weeks of age. We performed all animal work in accordance with reviewed/approved protocol by the Mayo Clinic Institutional Animal Care and Use Committee.

*CD4$^{+}$ T Cell Isolation from Human Intestinal Biopsy Specimens*

As previously described [23], human tissue biopsy specimens derived from the terminal ileum were rinsed in 0.9% NaCl and then placed in 1 mmol/L EDTA Duchmann media (RPMI 1640 supplemented with 10% fetal calf serum [FCS], 1% penicillin/streptomycin, 1% gentamicin sulfate, 10 mmol/L HEPES, and β-mercaptoethanol) for 15 minutes in a 37° C $CO_2$ incubator to remove epithelial cells. Biopsy specimens were digested in a 37° C $CO_2$ incubator with a cocktail of enzymes containing 1 mg/mL each of collagenase, DNase, and trypsin inhibitor for 1–2 h with gentle agitation. Digested tissue was filtered through a 70-µm cell strainer to obtain single cells, then centrifuged at 1200 rpm for 10 minutes. Cell pellet was subjected to CD4$^{+}$ T cell isolation according to the manufacturer's instructions (Miltenyi Biotec, San Diego, CA). Isolated CD4$^{+}$ T cells were subsequently subjected to *in situ* proximity ligation assay (PLA) to quantity mitochondria-ER and TOM20-IP3R1 interactions.

*CD4$^{+}$ T cell isolation from human PBMCs and mouse spleen, cell culture and differentiation*

Naive $CD4^+$ T cells were isolated from male mouse spleen ($CD62L^+$) or human PBMCs ($CD45RA^+$) using corresponding $CD4^+$ T cell isolation kit according to manufacturer's instructions (Miltenyi Biotec). All primary human and mouse $CD4^+$ T cells were cultured under standard conditions with cRPMI 1640 (RPMI 1640 + L-Glutamine [Gibco] supplemented with 10% FCS, 10 mmol/L HEPES pH 7.4, 1 mmol/L sodium pyruvate, 2 mmol/L L-glutamine, 1% non-essential amino acids, penicillin, streptomycin, and 50 µmol/L β-mercaptoethanol). Human $CD4^+$ T cell subsets were polarized from naïve cells in the presence of anti-human CD3 (2 µg/ml) and CD28 (2 µg/ml), and human IL-12 (20 ng/ml) for Th1 cells, human IL-4 (10 ng/ml), IL-2 (100 U/ml), and anti-human IFN-γ (10 µg/ml) for Th2 cells, human TGF-β1 (1.5 ng/ml), IL-1β (20 ng/ml), IL-6 (50 ng/ml), IL-21 (100 ng/ml), IL-23 (20 ng/ml), and anti-human IL-4 (2.5 µg/ml) and IFN-γ (5 µg/ml) antibodies for Th17 cells, and human TGF-β1 (5 ng/ml) and IL-2 (100 U/ml) for iTregs for 5 days with fresh media and cytokines added on day 2 and day 4. On day 5, differentiated T cells were harvested, resuspended in fresh media only and plated for 2-3 days in the presence of anti-human CD3 (2 µg/ml) and CD28 (2 µg/ml) antibodies. Differentiated (day 5) and developed (day 7-8) human $CD4^+$ T subsets were treated with indicated inhibitors or compounds in regular cRPMI or RPMI medium for the indicated time-point. $CD4^+CD25^+$ regulatory T cell isolation kit II (cat. 130-094-775; Miltenyi Biotec) was used to isolate natural Tregs (nTregs) from human PBMCs according to the manufacturer's instructions.

*Colitis Induction*

Male C57BL/6J mice and C57BL/6J *Rag1*$^{-/-}$ mice (Jackson Laboratory) were placed in conventional housing in the Mayo Clinic animal facility. For $CD4^+$ $CD45RB^{high}$ T cell-induced colitis induction, $CD4^+$ T cells were isolated from splenocytes of C57BL/6J mice. $CD4^+$ T cells were co-stained with CD45RB and CD25 fluorescently conjugated antibodies and sorted for $CD4^+$ $CD45RB^{high}$ T cells and $CD4^+$ $CD25^{++}$ (Tregs) via flow cytometry. 500,000 $CD4^+$ $CD45RB^{high}$ T cells were injected into *Rag1*$^{-/-}$ mice intraperitoneally (I.P.) along with vehicle control (PBS) or 250,000 WT Tregs or *Il21r*$^{-/-}$ Tregs (on day 0 for colitis prevention or on day 21 for colitis rescue). Weights of experimental mice were monitored for

0-7 weeks (disease rescue), or 0-12 weeks (disease prevention) while being fed non-irradiated chow. Wild-type and *Il21r*$^{-/-}$ (*Il21r*$^{tm1kopf}$) animals [22] (Jackson Laboratory) were housed in the Mayo Clinic animal facility for breading. Mice used in experiments were males of 6–9 weeks of age. All animal work was done in accordance with and reviewed/approved by the Mayo Clinic Institutional Animal Care and Use Committee.

*Colitis Assessment and Histopathology*

Mice were weighed every week and their colon lengths were determined during necropsy. The degree of colitis was quantified using three outcome variables: weight loss, colon histology, and a disease activity index. Mouse Colon Histology Index (MCHI) assesses eight parameters, including goblet cell loss, crypt density, crypt hyperplasia, muscle thickening, crypt abscess, ulceration, the extent of inflammatory infiltrate, and semiquantitative assessment of submucosal inflammation. The Disease Activity Index (DAI) is an established clinical index of colitis severity encompassing clinical signs of colitis (wasting and hunching of the recipient mouse and the physical characteristics of stool) and an ordinal scale of colonic involvement (thickness and erythema). Dissected colon was fixed in 10% neutral buffered formalin for 24 hours and embedded in paraffin. Tissue sections (4 µm) were cut using a microtome and positioned on glass slides. Hematoxylin and eosin (H&E) staining were performed according to standard techniques. H&E slides were reviewed by a gastrointestinal pathologist.

*Serum Cytokine Analysis*

Cytokine levels were determined in supernatants (50 µl of serum) using the BD cytometric bead array mouse Th1/ Th2/Th17 kit (BD Bioscience) according to the instructions of the manufacturer and analyzed using FCAP Array version 3 software (Soft Flow Hungary Ltd., Pécs, Hungary).

*Transmission Electron Microscopy*

Cells were harvested and washed with PBS. Cells were placed in fixative (4% paraformaldehyde + 1% glutaraldehyde in 0.1 M phosphate buffered saline, pH 7.2 (PBS). After fixation, cells were washed with PBS, suspended in 2% low melt agar, and spun down to the pellet. The agar suspended cells were stained with 1% osmium tetroxide, washed in $H_2O$, stained in 2% uranyl acetate, washed in $H_2O$, dehydrated through a graded series of ethanol and acetone, and embedded in Embed 812 resin. Following a 24 h polymerization at 60°C, 0.1 μM ultrathin sections were prepared and post-stained with lead citrate. Micrographs were acquired using a JEOL 1400 Plus transmission electron microscope (JEOL, Inc., Peabody, MA) at 80 kV equipped with a Gatan Orius camera (Gatan, Inc., Warrendale, PA). ImageJ software (NIH, Bethesda, USA) was used for measuring mitochondrial length (major axis) and width (minor axis). For morphological evaluation, the average aspect ratio was calculated using major axis to minor axis. An aspect ratio of around 1.5 indicates a circular mitochondrial section. To quantify mitochondria-ER contacts for each cell, total mitochondria, and mitochondria in contact with ER, were counted and the frequency of mitochondria-ER contacts calculated. Mitochondria-ER contacts were defined as mitochondria juxtaposed to ER at a distance of < 100 nm.

*Intracellular ATP measurement*

Cell lysates were subjected to ATP detection using manufacturer's instructions (ENLITEN ATP Assay System Bioluminescence Detection Kit for ATP Measurement from Promega, Cat #: FF2000), followed by measurement with Synergy H1-Multi Mode plate reader (BioTek, Winooski, VT).

*Serial Block-face Scanning Electron Microscopy (SEM)*

Samples were stained and prepared for serial block-face microscopy using an adapted protocol [24]. Briefly, tissue or cell samples were fixed by immersion in 2% glutaraldehyde + 2% paraformaldehyde in 0.15 M cacodylate buffer containing 2 mM calcium chloride until further processed (minimum of 24 h). Fixed samples were washed in 0.15 M cacodylate buffer and incubated at room temperature in 2% osmium tetroxide in 0.15 M cacodylate for 1.5 h. Without rinsing, samples were incubated in 2.5% potassium ferrocyanide + 2% osmium tetroxide in 0.15 M cacodylate for another 1.5 h. at room temperature. Following a rinse in $H_2O$, samples were incubated in 1% thiocarbohydrazide in $H_2O$ for 45 min at 50˚C. After another rinse in $H_2O$, samples were incubated sequentially in 2% osmium tetroxide in $H_2O$ for 1.5 h. at room temperature, 1% aqueous uranyl acetate overnight at 4˚C, and 7% lead aspartate solution 1 h at 50˚C, with several rinses in $H_2O$ between each reagent. Following dehydration through a series of ethanol and acetone, samples were infiltrated and eventually embedded in a hard formulation of Embed 812 resin (EMS, Hatfield, PA) and polymerized in a 60˚C oven for a minimum of 24 h. To prepare embedded samples for placement into the SEM and subsequent imaging, 1 $mm^3$ pieces were roughly trimmed of any excess resin and mounted to 8 mm aluminum stubs using silver epoxy Epo-Tek (EMS, Hatfield, PA). The mounted sample was then carefully trimmed to a 0.5 mm x 0.5 mm x 1 mm tall tower using a diamond trimming knife (Diatome Trimtool 45, EMS, Hatfield, PA). Trimmed sample and entire stub were coated with gold- palladium to assist in charge dissipation. The coated sample was then inserted into a VolumeScope serial block-face SEM (Thermo Fisher, Waltham, MA) and allowed to acclimate to high vacuum for 12 h prior to the start of imaging.

*Acquisition and post-processing of images*

High-resolution block-face images were obtained in a low vacuum environment using beam energy of 3.0 kV with a current of 100 pA and a scanning dwell time of 2 µs and a 10 nm pixel size. A stack of approximately 500 block-face images were obtained while cutting the block at 50 nm increments. The image stack was then aligned, filtered, segmented, and 3D rendering using Amira software (Thermo Fisher, Waltham, MA) with further analysis performed using Reconstruct [25].

*Proximity Ligation Assay (PLA) and Confocal Microscopy of Fixed Cells*

As previously described [23], cells were harvested and plated on 8-well Lab-Tek chamber slides (Thermo Fisher Scientific, Waltham, MA) coated with fibronectin (1 mg/ml) (Corning, Corning, NY) for 3 hours to allow cell attachment, and then stimulated or treated as indicated. Cells then were fixed with 4% paraformaldehyde, permeabilized with 0.15% Triton X-100 (Invitrogen, Carlsbad, CA), and washed with PBS. Cells were blocked for 1 h with 5% bovine serum albumin containing 0.1% glycine and incubated with anti-VDAC1 and anti-IP3R1 antibodies to biochemically measure mitochondria-ER junctions of < 30 nm distance. Mitochondria-ER junctions were visualized by Duolink in situ fluorescence PLA probes and detection reagents in accordance with the manufacturer's instructions (Sigma, St. Louis, MO). Cells were mounted with Duolink 40,6-diamidino-2-phenylindole–containing mounting (DAPI) medium (Sigma-Aldrich). Images of cells were captured by using a C-Apochromat 63_ objective/1.20 W korrM27 of a fluorescent confocal microscope (LSM780 AxioObserver; Carl Zeiss, Jena, Germany).

Images were processed using the Zen lite 2012 software (Carl Zeiss). Quantification of detected PLA signals or dots were measured using ImageJ (National Institute of Health, Bethesda, MD).

*Live-cell Confocal Imaging of Dye-labeled Cells*

As previously described [26], cells were stimulated as indicated in cRPMI media and plated on fibronectin-coated dishes for 2-3 h. Cells were stained with organelle tracking dyes (200-500 nM range) then imaged on an LSM 5 Live laser-scanning confocal microscope. Images of cells were captured using a C-Apochromat 63_ objective/1.20 W korrM27 of a fluorescent confocal microscope (LSM780 AxioObserver; Carl Zeiss, Jena, Germany). Images were processed using the Zen lite 2012 software (Carl Zeiss). Images were processed using the Zen lite 2012 software.

*Extracellular Flux Analysis using Agilent Seahorse XF Analyzer*

XFe-96 and XF-24 Extracellular Flux Analyzers (Seahorse Bioscience, Agilent Technologies) were used to define OCRs and ECARs. 300,000 – 500,000 cells were seeded in XF24 and XF96 Cell Culture Microplates before perturbation experiments. Mitochondrial perturbation experiments with cells were conducted in XF media containing glucose (25 mM or as otherwise indicated), pyruvate (2 mM) and L-glutamine (0.5 mM) by sequential addition of 1 μM oligomycin (Sigma-Aldrich), 1 μM FCCP (carbonyl cyanide 4-(trifluoromethoxy) phenylhydrazone; Sigma-Aldrich) and 0.5 μM rotenone/antimycin A (Sigma-Aldrich). Glycolysis stress tests with cells were conducted in XF media by sequential addition of 10 mM glucose (Sigma-Aldrich), 2 μM oligomycin (Sigma-Aldrich), and 50 mM 2-DG (Sigma-Aldrich). Changes in OCR and ECAR in cells were measured as indicated.

*Immunoblotting Analysis*

Cells were harvested, washed with cold PBS, and lysed with Radioimmunoprecipitation (RIPA) lysis buffer containing 50 mmol/L Tris-HCl, pH 7.4, 1% NP-40, 150 mmol/L NaCl, 2 mmol/L EDTA, and protease and phosphatase inhibitors.  Cell lysates were boiled in SDS sample buffer for 10 mins at 95-100° C and resolved on SDS-PAGE gel and transferred to PVDF membranes. Membranes were blocked for 1 h in milk or BSA and then incubated with corresponding primary antibodies at 4°C overnight. After washing, membranes were incubated with secondary antibodies in TBS + Tween 20 +

5% nonfat dry milk for 1 h. ECL western blotting chemiluminescent substrates were added for 1 min and membranes were imaged on Bio-Rad imaging software. Membranes were stripped and re-probed with antibodies.

*Mass Cytometry by Time-of-Flight (CyTOF)*

High-dimensional CyTOF was utilized to examine the expression of human iTreg surface markers and intracellular cytokines under varying conditions in the absence of brefeldin. Cells were individually barcoded with a unique combination of palladium isotopes (Fluidigm, Cat#201060). All samples were then pooled and stained with a custom panel of heavy metal conjugated antibodies (see Supplementary Table1 in supplementary Methods for list of antibodies), and data was obtained on a Helios mass cytometer (Fluidigm). Data cleanup, de-barcoding, and quality control was performed by the Mayo Clinic Immune Monitoring Core. Data analysis was performed primarily using CytoBank by gating on live cells, singlets, $CD45^+$, $CD3^+$, $CD4^+$, $CD127^-$ cells. iTregs highly co-expressing CD25 and FOXP3 (80% of cells) were gated on and examined for changes in protein expression. Median protein expression values were performed using GraphPad Prism. The median expression of protein is indicated in the heat map.

*Extracellular and Intracellular Metabolomics*

For Gas chromatography-mass spectrometry (GC-MS) analysis, methods are described as before [27]. Briefly, 3 million iTreg cells stimulated as indicated in normal culture medium were quickly removed, cells were washed 3 times by saline, quenched with -20°C 1:1 water: methanol, and flash frozen in liquid nitrogen. Cells were thawed on ice for 20 minutes. After vortexing, samples were centrifuged at 10,000 g for 10 minutes at 4 °C. The supernatant was then collected and completely dried in a SpeedVac concentrator, subsequently methoximated using 20 µL MOX™ Regent at 30°C for 90 min and then derivatized using 80 µL of MSTFA+1% TMCS (N-methyl-N-trimethylsilyltrifluoroacetamide with 1% trimethylchlorosilane) at 37°C for 30 min. Metabolite levels were determined using GC-MS (Hewlett-Packard, HP 5980B) with DB5-MS column. GC-MS spectra were deconvoluted using AMDIS software, after that SpectConnect software was used to create metabolite peaks matrix. The Agilent

Fiehn GC/MS Metabolomics RTL Library was used for metabolite identifications. Ion count peak area was used for analysis of the relative abundance of the metabolites [28, 29]. For Nuclear Magnetic Resonance (NMR) spectroscopy analysis, protein from cell culture medium (300 µL) was precipitated by adding 900 µL ice-cold methanol. After vortexing, samples were centrifuged at 10,000 g for 10 minutes at 4 °C. The supernatant was then collected and completely dried in a SpeedVac concentrator. Samples were reconstructed by adding 500 µL of 0.1M phosphate buffer and 50 µL of 1 mM TSP-d4 solution in $D_2O$. Samples were vortexed for 20 seconds and transferred to 5 mm NMR tubes. NMR spectra were acquired on a Bruker 500 MHz Avance III HD spectrometer equipped with a BBO cryoprobe and SampleCase auto sampler (Bruker Biospin, Rheinstetten, Germany). $^1$H-NMR spectra were recorded using 1D noesy pulse sequence with presaturation (noesygppr1d), with 90-degree pulse (~13 µs), 4.68 seconds acquisition time, and 4 seconds relaxation delay. Spectra were phase and baseline corrected using the Topspin 3.5 software. Metabolites were identified and quantified using the software program Chenomx NMR Suite 8.2, by fitting the spectral lines of library compounds into the recorded NMR spectrum of the cell medium. The quantification was based on peak area of TSP-d4 signal, and metabolite concentrations were reported as µM in medium [27].

*Flow Cytometry*

Cells were treated as indicated with or without brefeldin A (BioLegend), fixed with True-Nuclear Transcription Factor Buffer Fix solution (BioLegend), permeabilized with Perm/Wash buffer (BioLegend), blocked for 10 minutes and then stained with relevant fluorochrome-conjugated primary antibodies. Cells were washed twice with Perm/Wash buffer and subsequently subjected to flow cytometry for fluorescence-activated cell sorting. Cells were electronically gated on live cells for analysis.

*RNA extraction and Metabolic Microarray Analysis*

Extracted RNA was subjected to mRNA analysis using the NuRNA™ Human Central Metabolism PCR Array (RNA ARRAYSTAR INC: Roche Light Cycler 480) according to manufacturers' instructions. This systematically profiles the expression of 373 transcripts encoding the enzymes or proteins involved in cellular metabolism. Briefly, cDNA was obtained from RNA using rtSTAR™ First-Strand cDNA Synthesis Kit and subjected to Arraystar SYBR Green Real-time qPCR Master Mix.

*Real-time (RT) Quantitative PCR*

Total RNA was isolated with RNeasy Plus Mini Kit (Qiagen) and was reverse transcribed using iScript cDNA synthesis kit (Bio-Rad). Quantification of gene expression was performed by real-time polymerase chain reaction using SYBR green fluorescence on a LightCycler 480 instrument (Roche). Specific primers are listed in Supplementary Table 2 in Supplementary Methods. Target gene expression was calculated using ΔΔCt method. Expression was normalized to 18S expression levels, which were stable across all experimental groups.

*SiRNA-mediated Gene Silencing*

For siRNA transfection, we used the smartpool Accell Human VDAC1 siRNA SMARTPool (Cat ID: E-019764-00-0050) and Accell Non-Targeting Control Pool (Cat ID:D-001910-10-50) according to manufacturer's (Dharmacon™) recommendations with minor modifications. Human iTregs were cultured in RPMI media supplemented with 2.5% FCS containing 5 µM siRNA for 4 days (day 5 – day 9) in the presence of anti-CD3 and anti-CD28 antibodies.

*Transcriptome Analysis*

Previously published single-cell RNA-seq data generated based on uninflamed and inflamed ileal resection specimens isolated from Crohn's disease patients were analyzed as described earlier [20]. Briefly, sequencing data were aligned to the human reference genome Grch38. Data with at least 500 unique molecular identifiers (UMIs), log10 genes per UMI >0.8, >250 genes per cell and a mitochondrial ratio of less than 0.2% were extracted, normalized, and integrated using the seurat package v3.0 in R 4.0.2. Tregs (*FOXP3*$^+$, *CD4*$^+$, *IL2RA*$^+$) were compared between inflamed and uninflamed samples as well as inflamed resection specimens of GIMATS and non-GIMATS data sets. Statistical differences were determined using unpaired t-test.

*Statistical Analysis*

Please refer to the Fig legends for description of sample size (*n*) and statistical details. Two-tailed unpaired Student's t test was used to compare two groups. One-way or two-way ANOVA + Dunn's or Bonferroni multiple comparisons test was used to compare three or more groups. Non-parametric murine data (DAI and MCHI) were analyzed using Kruskal-Wallis test or two-tailed unpaired Student's t test. *P* values less than 0.05 were considered statically significant. All data were analyzed using Prism (GraphPad Software, San Diego, CA).

*Data Availability*

The microarray data generated in this study has been deposited in the GEO database under accession code GSE224033. All other data supporting our findings are available within the article and its supplementary files.

Supplementary Table 1**:** Panel of extracellular markers and intracellular cytokines *metal*-isotope-tagged antibodies utilized for CyTOF analysis.

| *Protein Target* | *Antibody Clone* | *Antibody Tag* | *Manufacturer* |
|---|---|---|---|
| CD49D | 9F10 | 141Pr | Fluidigm |
| CD4 | RPA-T4 | 145Nd | Fluidigm |
| CCR4 | L291H4 | 149Sm | Fluidigm |
| CD45RA | HI100 | 153Eu | Fluidigm |
| CD3 | UCHT1 | 154Sm | Fluidigm |
| CD39 | A1 | 160Gd | Fluidigm |
| FOXP3 | PCH101 | 162Dy | Fluidigm |
| CD95 | DX2 | 164Dy | Fluidigm |
| CD45RO | UCHL1 | 165Ho | Fluidigm |
| CD25 | 2A3 | 169Tm | Fluidigm |
| CD152 | 14D3 | 170Er | Fluidigm |
| HLA-DR | L243 | 174Yb | Fluidigm |
| CD127 | A019D5 | 176Yb | Fluidigm |
| IL-5 | TRFK5 | 151Eu | Fluidigm |
| TNF-α | Mab11 | 152Sm | Fluidigm |
| TGF-β1 | TW4-6H10 | 163Dy | Fluidigm |
| IL-10 | JES3-9D7 | 166Er | Fluidigm |
| IL-17A | N49-653 | 172Yb | Fluidigm |
| IFN-γ | B27 | 168Er | Fluidigm |

Supplementary Table 2: Primer sets for RT-qPCR

| *GENE NAMES* | *HUMAN GENE PRIMER SETS (5'-3')* |
|---|---|
| *HIF1A* Forward | GAACGTCGAAAAGAAAAGTCTCG |
| *HIF1A* Reverse | CCTTATCAAGATGCGAACTCACA |
| *LDHA* Forward | AGCTGTTCCACTTAAGGCCC |
| *LDHA* Reverse | TGGAACCAAAAGGAATCGGGA |
| *CS* Forward | TGCTTCCTCCACGAATTTGAAA |
| *CS* Reverse | CCACCATACATCATGTCCACAG |
| *GLUT1* Forward | ATTGGCTCCGGTATCGTCAAC |
| *GLUT1* Reverse | GCTCAGATAGGACATCCAGGGTA |
| *GLUT3* Forward | GCTCTCTGGGATCAATGCTGTGT |
| *GLUT3* Reverse | CTTCCTGCCCTTTCCACCAGA |
| *HK2* Forward | AACAGCCTGGACGAGAGCAT |
| *HK2* Reverse | GCCAACAATGAGGCCAACTT |
| *GPI* Forward | GATGGTAGCTCTCTGCAGCC |
| *GPI* Reverse | GCCATGGCGGGACTCTTG |
| *PFK* Forward | GGCAGCCATGCATAAAGACG |
| *PFK* Reverse | AAGCTTCCCCAGCTGTTCTC |
| *TPI* Forward | AGGCATGTCTTTGGGGAGTC |
| *TPI* Reverse | AGTCCTTCACGTTATCTGCGA |
| *ENO1* Forward | CGCCTTAGCTAGGCAGGAAG |
| *ENO1* Reverse | GGTGAACTTCTAGCCACTGGG |
| *PKM2* Forward | ACGAGAACATCCTGTGGCTG |
| *PKM2* Reverse | AGGAAGTCGGCACCTTTCTG |
| *CPT1* Forward | ATCAATCGGACTCTGGAAACGG |

| | |
|---|---|
| *CPT1* Reverse | TCAGGGAGTAGCGCATGGT |
| *ACLY* Forward | ATCGGTTCAAGTATGCTCGGG |
| *ACLY* Reverse | GACCAAGTTTTCCACGACGTT |
| *ACC1* Forward | ATGTCTGGCTTGCACCTAGTA |
| *ACC1* Reverse | CCCCAAAGCGAGTAACAAATTCT |
| *ACC2* Forward | CAA GCCCATCACCAAGAGTAAA |
| *ACC2* Reverse | CCCTGAGTTATCAGAGGCTGG |
| *FASN* Forward | ACAGCGGGGAATGGGTACT |
| *FASN* Reverse | GACTGGTACAACGAGCGGAT |
| *HMGCR* Forward | GTGAGATCTGGAGGATCCAAGG |
| *HMGCR* Reverse | GATGGGAGGCCACAAAGAGG |
| *HMGCS1* Forward | GTTGGCGGCTATAAAGCTGGT |
| *HMGCS1* Reverse | CCTTCGGGCACAAGCG |
| *GAPDH* Forward | AGCCGCATCTTCTTTTGCGTCG |
| *GAPDH* Reverse | GACCAGGCGCCCAATACG |
| *FOXP3* Forward | CGGACCATCTTCTGGATGAG |
| *FOXP3* Reverse | TTGTCGGATGATGCCACAG |
| *IL17A* Forward | GGATGTTCAGGTTGACCATCAC |
| *IL17A* Reverse | TCCCACGAAATCCAGGATGC |
| *IL17F* Forward | GCTGTCGATATTGGGGCTTG |
| *IL17F* Reverse | GGAAACGCGCTGGTTTTCAT |
| *TNFA* Forward | CCTCTCTCTAATCAGCCCTCTG |
| *TNFA* Reverse | GAGGACCTGGGAGTAGAG |
| *IFNG* Forward | TCGGTAACTGACTTGAATGTCCA |
| *IFNG* Reverse | TCGCTTCCCTGTTTTAGCTGC |

| | |
|---|---|
| *IL10* Forward | GACTTTAAGGGTTACCTGGGTTG |
| *IL10* Reverse | GATGTCAAACTCACTCATGGCT |
| *TGFB1* Forward | GGCCAGATCCTGTCCAAGC |
| *TGFB1* Reverse | GTGGGTTTCCACCATTAGCAC |
| *MDH1* Forward | GGTGCAGCCTTAGATAAATACGC |
| *MDH1* Reverse | AGTCAAGCAACTGAAGTTCTCC |
| *18S* Forward | CGCTTCCTTACCTGGTTGAT |
| *18S* Reverse | GAGCGACCAAAGGAACCATA |
| *VDAC1* Forward | ACGTATGCCGATCTTGGCAAA |
| *VDAC1* Reverse | TCAGGCCGTACTCAGTCCATC |

*Key Resources Table*

| REAGENT or RESOURCE | SOURCE | IDENTIFIER |
|---|---|---|
| *Antibodies* | | |
| Anti-human/mouse/rat VDAC1 (Lot #3507477001) | ORIGENE | Cat#TA326858 |
| Anti-human/mouse/rat/monkey GRP75 | Abcam | Cat#ab2799 |
| Anti-human/mouse/rat/monkey TOM20 | Cell Signaling | Cat#42406S |
| Anti-human/mouse/rat IP3R-I | Santa Cruz | Cat#sc-271197 |
| Alexa Fluor 488 Anti-human FOXP3 (clone 259D) | BioLegend | Cat#320212 |
| Alexa Fluor 488 Anti-human CD25 (clone BC96) | BioLegend | Cat#302616 |
| PE anti-human CD39 (clone A1) | BioLegend | Cat#328208 |
| PerCP/Cy5.5 anti-human CD152 (CTLA4) (clone BNI3) | BioLegend | Cat#369608 |
| Anti-human/mouse/rat/monkey phospho-Serine 9 GSK3β (D85E12) | Cell Signaling | Cat#5558S |
| Anti-human/mouse/rat/monkey GSK3β (27C10) | Cell Signaling | Cat#9315 |
| Anti-human/mouse/rat/monkey β-actin (8H10D10) | Cell Signaling | Cat#3700S |
| Anti-human/mouse/rat/monkey STAT3 (124H6) | Cell Signaling | Cat#9139S |
| Anti-human/mouse/rat HK-I | Invitrogen | Cat#MA5-15680 |
| APC anti-human CD49D (α4) (clone 9F10) | BioLegend | Cat#304308 |
| PE anti-human CD103 (integrin αE) (clone Ber-ACT8) | BioLegend | Cat#350206 |
| APC/Cyanine7 anti-human IFN-γ (4S.B3) | BioLegend | Cat#502530 |
| PE anti-human TNF-α (MAb11) | BioLegend | Cat#502909 |
| Alexa Fluor 488 anti-human CD4 (clone RPA-T4) | BioLegend | Cat#300519 |
| PE anti-mouse CD45RB (clone c363-16A) | BioLegend | Cat#103308 |

| | | |
|---|---|---|
| Alexa Fluor 488 CD25 (clone PC61) | BioLegend | Cat#102017 |
| Alexa Fluor 647 anti-human phospho-Serine 9 GSK3β | R&D SYSTEMS | Cat# IC25062R |
| Alexa Fluor 488 Mouse IgG1, k (clone MOPC-21) | BioLegend | Cat#400134 |
| APC Mouse IgG1, k (clone MOPC-21) | BioLegend | Cat#400119 |
| PE Mouse IgG1, k (clone MOPC-21) | BioLegend | Cat#400114 |
| APC/Cyanine 7 Mouse IgG1, k (clone MOPC-21) | BioLegend | Cat#400128 |
| Brilliant Violet 421 Mouse IgG1, k (clone MOPC-21) | BioLegend | Cat#400157 |
| PE/Cy7 Mouse IgG1, k (clone MOPC-21) | BioLegend | Cat#400125 |
| PercP/Cy5.5 Mouse IgG1, k (clone MOPC-21) | BioLegend | Cat#400150 |
| Human TruStain FcX™ | BioLegend | Cat#422302 |
| CellTak | Corning | Cat#354242 |
| *Chemicals, Peptides, and Recombinant Proteins* | | |
| LY2090314 | Selleckchem | Cat#S7063 |
| TCS 2002 | TOCRIS | Cat#3869 |
| Methyl Pyruvate | Sigma-Aldrich | Cat#371173 |
| MOXTM Regent | Thermo Fisher Scientific | Cat#TS-45950 |
| MSTFA+1% TMCS | Thermo Fisher Scientific | Cat#TS-48915 |
| Methanol CHROMASOLV®, for HPLC | Sigma-Aldrich | Cat#34860 |
| Water for HPLC | Sigma-Aldrich | Cat#270733 |
| TSP-d4 | Sigma-Aldrich | Cat#269913 |
| MitoTracker RedCMXRos | Thermo Fisher Scientific | Cat#M7512 |
| ER-Tracker™ Blue-White DPX | Thermo Fisher Scientific | Cat#E12353 |

| | | |
|---|---|---|
| Oligomycin | Abcam | Cat#ab141829 |
| FCCP | Abcam | Cat#ab120081 |
| Rotenone | Abcam | Cat#ab143135 |
| Antimycin A | Abcam | Cat#ab141904 |
| 2-deoxyglucose (2-DG) | APEXBIO | Cat#B1027 |
| Seahorse XF 1.0 M Glucose solution | Agilent Technologies | Cat#103577-100 |
| Seahorse XF 200 mM Glutamine solution | Agilent Technologies | Cat#103579-100 |
| Seahorse XF 100 mM Pyruvate solution | Agilent Technologies | Cat#103578-100 |
| XF RPMI Medium | Agilent Technologies | Cat#103576-100 |
| Purified NA/LE Mouse Anti-human CD3 | BD Pharmingen | Cat#555329 |
| Purified NA/LE Mouse Anti-human CD28 | BD Pharmingen | Cat#555725 |
| Recombinant human IL-2 | PetroTech | Cat#200-02 |
| Recombinant human TGF-β1 | PetroTech | Cat#100-21C |
| Recombinant human IL-12 p70 | PetroTech | Cat#200-12 |
| Recombinant human IL-1β | PetroTech | Cat#200-01B |
| Recombinant human IL-6 | PetroTech | Cat#200-06 |
| Recombinant human IL-21 | PetroTech | Cat#200-21 |
| Recombinant human IL-23 | PetroTech | Cat#200-23 |
| Recombinant human IL-4 | PetroTech | Cat#200-04 |
| Purified anti-human IFN-γ (clone B27) | BioLegend | Cat#506502 |
| Purified anti-human IL-4 (clone 8D4-8) | BioLegend | Cat#500702 |
| Clotrimazole | TOCRIS | Cat#4096 |
| Bifonazole | Sigma-Aldrich | Cat#B3563 |
| UK5099 | EMD Millipore | Cat#5.04817.0001 |
| TEPP-46 | Selleckchem | Cat#S7302 |

| | | |
|---|---|---|
| (ML265, CID-44246499, NCGC001186528) | | |
| BMS-303141 | Sigma-Aldrich | Cat#SML0784 |
| Cpd9, ACC1 inhibitor | EMD Millipore | Cat#5.34335.0001 |
| 2-Thenoyltrifluoroacetone (TTFA) | Sigma-Aldrich | Cat#T27006 |
| Sodium Fluoroacetate | MP, Biochemicals, LLC | Cat#201080 |
| Etomoxir sodium salt hydrate | Sigma-Aldrich | Cat#E1905 |
| Fibronectin Human, Natural | Corning | Cat#354008 |
| MitoTracker Green | Thermo Fisher Scientific | Cat#M7514 |
| Purified NA/LE Hamster Anti-mouse CD3e (clone145-2C11) | BD Pharmingen | Cat#553057 |
| Purified NA/LE Hamster Anti-mouse CD328 (clone145-2C11) | BD Pharmingen | Cat#553294 |
| Recombinant mouse IL-12 p70 | PetroTech | Cat#210-12 |
| Recombinant mouse IL-1β | PetroTech | Cat#211-11B |
| Recombinant mouse IL-6 | PetroTech | Cat#210-16 |
| Recombinant mouse IL-21 | PetroTech | Cat#210-21 |
| Recombinant mouse IL-23 p40 | R&D SYSTEMS | Cat#1887-ML-010 |
| LEAF$^{TM}$ Purified anti-mouse IFN-γ (clone XMG1.2) | BioLegend | Cat#505812 |
| Purified anti-mouse IL-4 (clone 11B11) | BioLegend | Cat#504102 |
| *Biological samples and Experimental Models* | | |
| Healthy adult – peripheral blood mononuclear cells (males) | Mayo Clinic | N/A |
| Healthy adult – intestinal biopsies (male) | Mayo Clinic | N/A |
| C57BL/6J mice (males) | The Jackson Laboratory | Stock no:000664 |
| C57BL/6J *Rag1*$^{-/-}$ mice (males) | The Jackson Laboratory | Stock no: 002216 |

| | | |
|---|---|---|
| C57BL/6NJ wildtype (one male and one female) | The Jackson Laboratory | Stock no: 005304 |
| Total body *Il21r*$^{-/-}$ (one male and one female) | The Jackson Laboratory | Stock no: 019115 |
| *Software and Algorithms* | | |
| FlowJo version10.2 | FlowJo, LLC | https://www.flowjo.com |
| Zen 3.0 (blue edition) | Zen | https://www.zeiss.com |
| Bio-Rad Image Lab | Bio-Rad | https://www.bio-rad.com |
| GraphPadPrism 8 | GraphPad software | https://www.graphpad.com |
| Seahorse Bioscience Wave Desktop | Agilent Technologies | https://www.agilent.com |
| ImageJ | NIH | https://imagej.nih.gov |
| Photoshop Adobe | Adobe | https://www.adobe.com |
| *Critical Commercial Assays* | | |
| BD™ Cytometric Bead Array (CBA) Mouse Th1/Th2/Th17 Cytokine Kit | BD Biosciences | Cat#560485 |
| ENLITEN ATP Assay System | Promega | Cat#FF2000 |
| Human Treg Phenotyping Panel Kit | Fluidigm | Cat#201060 |
| RNeasy Plus Mini Kit | QIAGEN | Cat#74134 |
| Seahorse XF24 Flux/Pak | Agilent Technologies | Cat#100850-001 |
| Seahorse XFe96 Flux/Pak | Agilent Technologies | Cat#102416-100 |
| Arraystar SYBR Green qPCR Mater Mix | ARRAYSTAR INC | Cat#AS-MR-006-5 |
| NuRNA™ Human Central Metabolism PCR Array (Roche Light Cycler 480) | ARRAYSTAR INC | Cat#AS-NM-004-1-R |
| Spike-in RNA | ARRAYSTAR INC | Cat#AS-SP-001-1 |
| Duolink™ In Situ PLA Anti-Mouse PLUS | Sigma-Aldrich | Cat#DUO92001 |
| Duolink™ In Situ PLA Anti-Rabbit MINUS | Sigma-Aldrich | Cat#DUO92005 |
| Duolink™ In Situ Detection Reagents Red | Sigma-Aldrich | Cat#DUO92008 |

| | | |
|---|---|---|
| $CD4^+$ T cell isolation kit, mouse | Miltenyi Biotec | Cat#130-104-454 |
| $CD4^+$ $CD62L^+$ (naïve) T cell isolation kit, mouse | Miltenyi Biotec | Cat#130-106-643 |
| $CD4^+$ T cell isolation kit, human | Miltenyi Biotec | Cat#130-096-533 |
| Accell Human VDAC1 siRNA SMARTPool | Dharmacon™ | Cat ID: E-019764-00-0050 |
| Accell Non-targeting Control Pool | Dharmacon™ | Cat ID: D-001910-10-50 |
| $CD45RA^+$ (naïve) cell isolation kit, human | Miltenyi Biotec | Cat#130-045-901 |
| $CD4^+$ $CD127^{dim}$ T cell isolation kit, human | Miltenyi Biotec | Cat#130-094-775 |
| *Others* | | |
| RPMI Medium 1640 (1X) [+] L-Glutamine | Gibco | REF#11875-093 |
