## Supplementary figures for "Interleukin-21 Drives a Hypermetabolic State and CD4^+^ T Cell-associated Pathogenicity in Chronic Intestinal Inflammation"

**Short title**: IL-21 induces metabolic disturbance in Tregs

Adebowale O. Bamidele[1, 2], Shravan K. Mishra[1], Petra Hirsova[3], Patrick J. Fehrenbach[1], Lucia Valenzuela-Pérez[3], Hyun Se Kim Lee[3]

[1]Immunometabolism and Mucosal Immunity Laboratory, Division of Gastroenterology and Hepatology, Mayo Clinic, 200 First Street SW, Rochester, MN 55905, USA
[2]Department of Immunology, Mayo Clinic, 200 First Street SW, Rochester, MN 55905, USA
[3]Division of Gastroenterology and Hepatology,
Mayo Clinic, 200 First Street SW, Rochester, MN 55905, USA

### Supplementary Figures and Legends

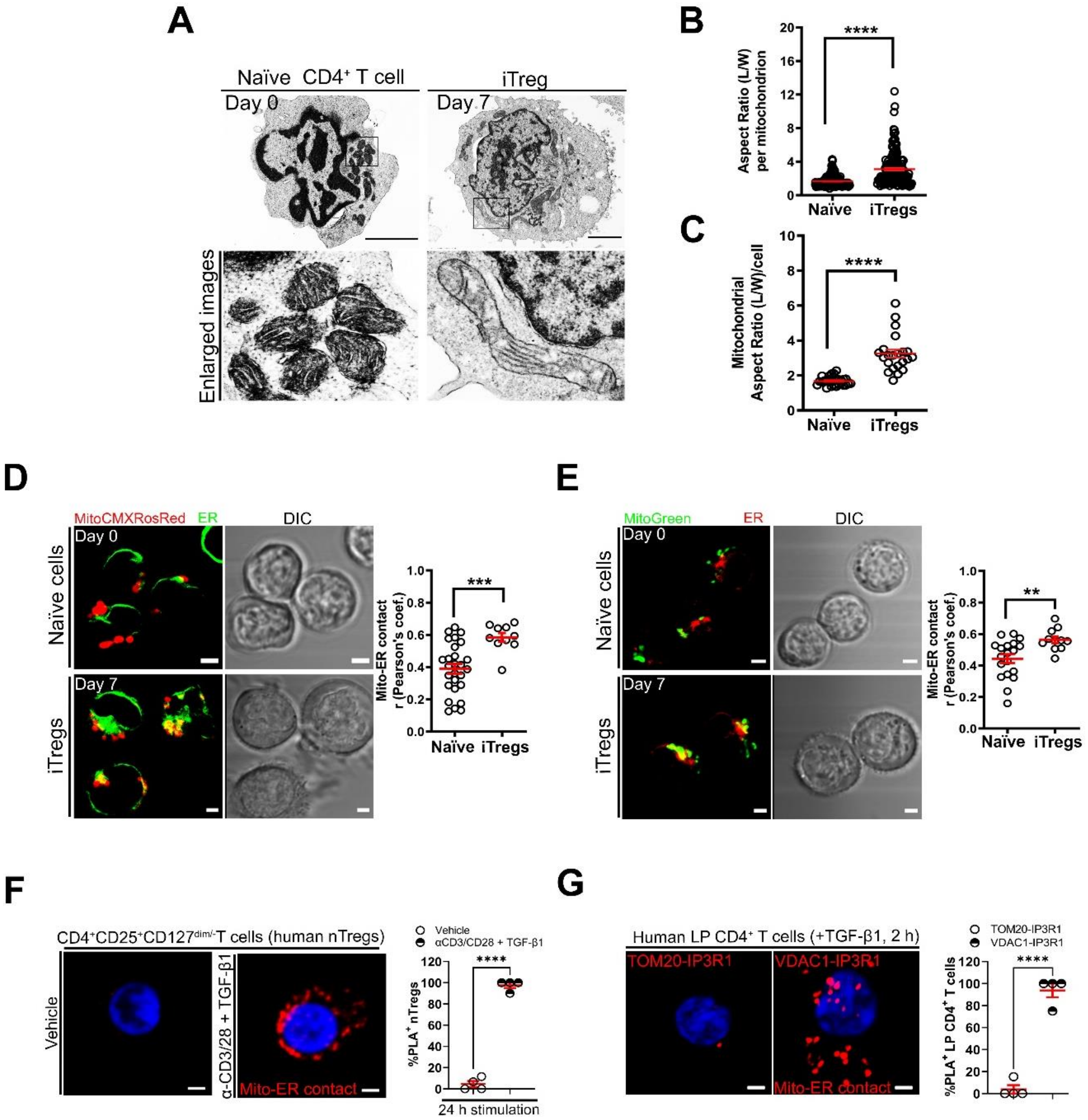


**Supplementary Figure 1. Mitochondria-ER Appositions are Present in Human Tregs.**

(A-C) Representative TEM images with enlarged images of naïve cells (n = 29) and iTregs (n = 22) from n = 3 biological replicates; scale bar, 5 µm (A). Graphs show aspect ratio per mitochondrion (n = 252 vs. n = 162 mitochondria) (B) and mitochondrial aspect ratio per cell (n = 29 vs. n = 22) (C).

(D and E) Representative confocal images of live cells loaded with organelle tracker dyes (MitoTracker Red chloromethyl-X-rosamine, CMX-Ros; Blue-White DPX, ER-Tracker in green pseudo color (D) (n =

27 vs. n = 10); MitoTracker Green, Blue-White DPX, ER-Tracker in red pseudo color (E) (n = 19 vs. n = 11). Graphs show mito-ER co-localization measured by the Pearson's correlation coefficient per cell (Pearson's coef.); scale bars, 5 μm.

(F) Representative PLA images show mito-ER contact in red in activated and TGF-β1-stimulated human nTregs (24 h) (n = 4); scale bar, 2 μm. Graph shows PLA signals (n = 56 vs. n = 82).

(G) Representative PLA images show VDAC1-IP3R1 binding vs. TOM20-IP3R1 binding in TGF-β1-stimulated LP $CD4^+$ T cells (25 ng/ml, 2 h) (n = 2); scale bar, 2 μm. Graph shows PLA signals (n = 35 vs. n = 29).

Data represents mean ± SEM. * $p < 0.05$, ** $p < 0.01$, *** $p < 0.001$, and **** $p < 0.0001$, using two-tailed Student's t test.

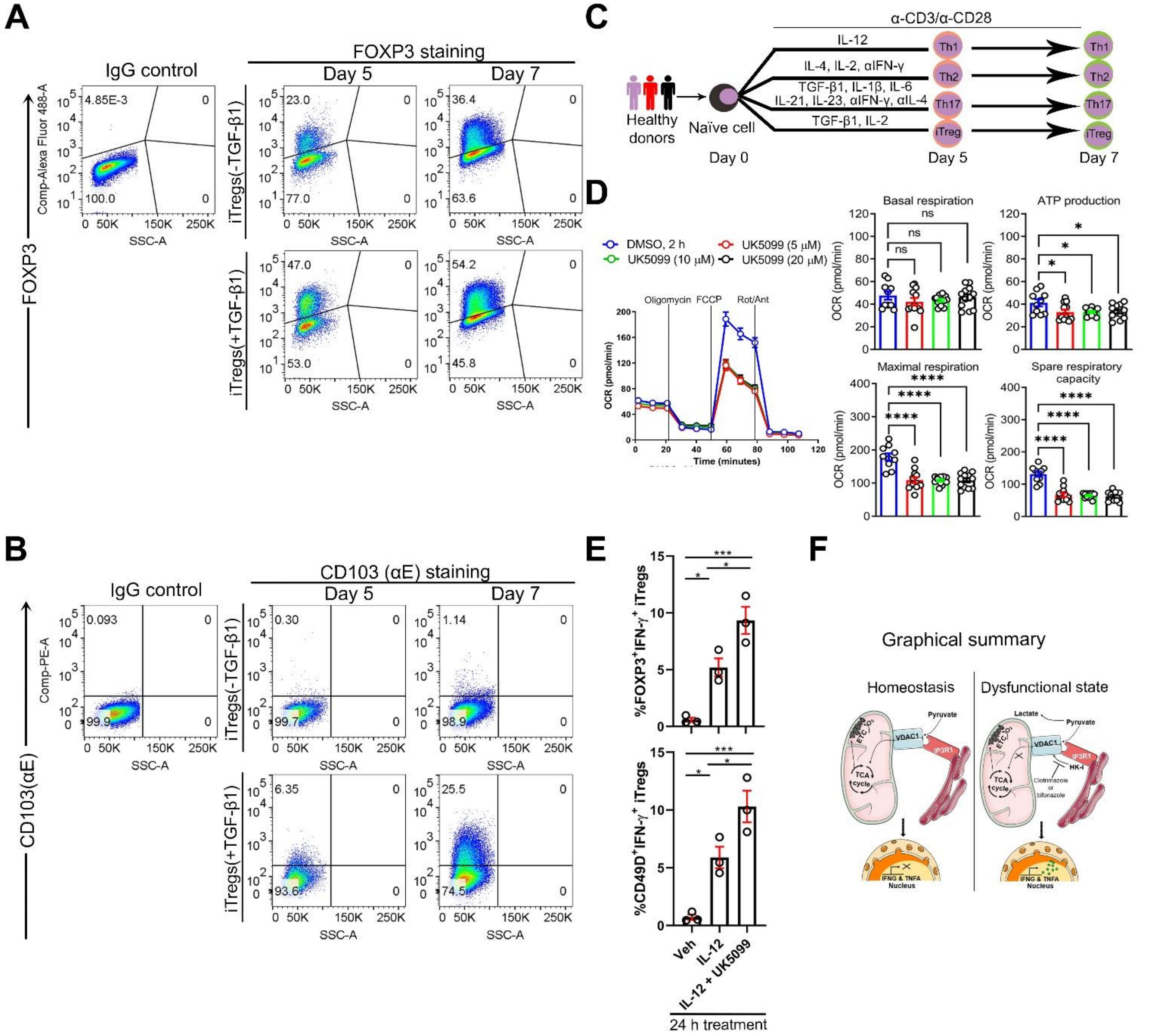


**Supplementary Figure 2. VDAC1 Inhibition Alters iTreg Metabolic State and Sensitizes to IL-12-induced Inflammatory Response.**

(A and B) Dot plots show the percentage of FOXP3$^+$ (A) or CD103$^+$ (B) iTregs.

(C) Experimental workflow for the differentiation of human naïve cells into CD4$^+$ T cell subsets.

(D) Representative OCR profile of iTregs ± UK5099 (n = 3). Bar graphs show calculated basal respiration, ATP production, maximal respiration, and spare respiratory capacity; mean ± SEM from 10-12 technical replicates.

(E) Percentage of FOXP3$^+$ IFN-γ$^+$ (top) or CD49D(α4)$^+$ IFN-γ$^+$ (bottom) iTregs ± IL-12 (25 ng/ml), UK5099.

(F) Graphical summary for the role of pyruvate transport in preventing iTreg inflammatory response.

Data represents mean ± SEM. * $p < 0.05$, *** $p < 0.001$, and **** $p < 0.0001$, using two-tailed Student's t test or one-way ANOVA followed by Bonferroni test for multiple comparisons.

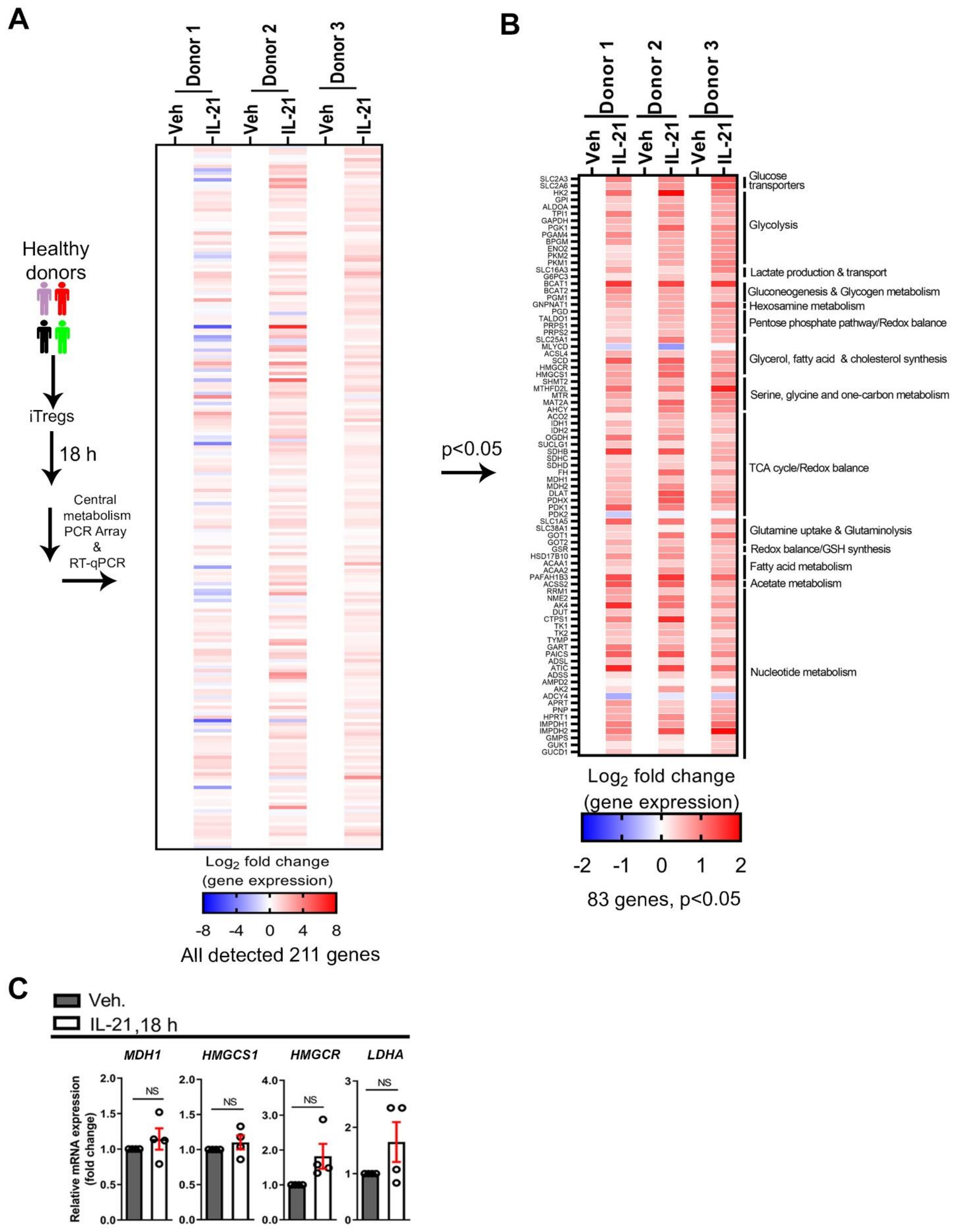
A
Healthy donors
iTregs
18 h
Central metabolism PCR Array & RT-qPCR
Donor 1
Donor 2
Donor 3
Veh
IL-21
Log$_2$ fold change (gene expression)
-8 -4 0 4 8
All detected 211 genes
p<0.05
B
SLC2A3
SLC2A6
HK2
GPI
ALDOA
TPI1
GAPDH
PGK1
PGAM4
BPGM
ENO2
PKM2
PKM1
SLC16A3
G6PC3
BCAT1
BCAT2
PGM1
GNPNAT1
PGD
TALDO1
PRPS1
PRPS2
SLC25A1
MLYCD
ACSL4
SCD
HMGCR
HMGCS1
SHMT2
MTHFD2L
MTR
MAT2A
AHCY
ACO2
IDH1
IDH2
OGDH
SUCLG1
SDHB
SDHC
SDHD
FH
MDH1
MDH2
DLAT
PDHX
PDK1
PDK2
SLC1A5
SLC38A1
GOT1
GOT2
GSR
HSD17B10
ACAA1
ACAA2
PAFAH1B3
ACSS2
RRM1
NME2
AK4
DUT
CTPS1
TK1
TK2
TYMP
GART
PAICS
ADSL
ATIC
ADSS
AMPD2
AK2
ADCY4
APRT
PNP
HPRT1
IMPDH1
IMPDH2
GMPS
GUK1
GUCD1
Glucose transporters
Glycolysis
Lactate production & transport
Gluconeogenesis & Glycogen metabolism
Hexosamine metabolism
Pentose phosphate pathway/Redox balance
Glycerol, fatty acid & cholesterol synthesis
Serine, glycine and one-carbon metabolism
TCA cycle/Redox balance
Glutamine uptake & Glutaminolysis
Redox balance/GSH synthesis
Fatty acid metabolism
Acetate metabolism
Nucleotide metabolism
Log$_2$ fold change (gene expression)
-2 -1 0 1 2
83 genes, p<0.05
C
Veh.
IL-21,18 h
MDH1
HMGCS1
HMGCR
LDHA
Relative mRNA expression (fold change)
NS

**Supplementary Figure 3. IL-21 Stimulation of Human iTregs Promotes a Hypermetabolic State.**

(A) Heatmap shows all detected 211 metabolic genes (IL-21/vehicle) in iTregs.

(B) Heatmap of 83 relevant metabolic transcripts (IL-21/vehicle, $p < 0.05$) and associated metabolic processes.

(C) Relative mRNA expression via RT-qPCR of enzymes associated with mitochondrial citrate synthesis (malate dehydrogenase; MDH1), cholesterol biosynthesis (3-hydroxy-3-methyl-glutaryl-CoA synthase 1; HMGCS1 and 3-hydroxy-3-methyl-glutaryl-CoA reductase; HMGCR), and lactate production (lactate dehydrogenase; LDHA) in iTregs ± IL-21; mean ± SEM from n = 4, Mann-Whitney *U* test, NS (not significant).

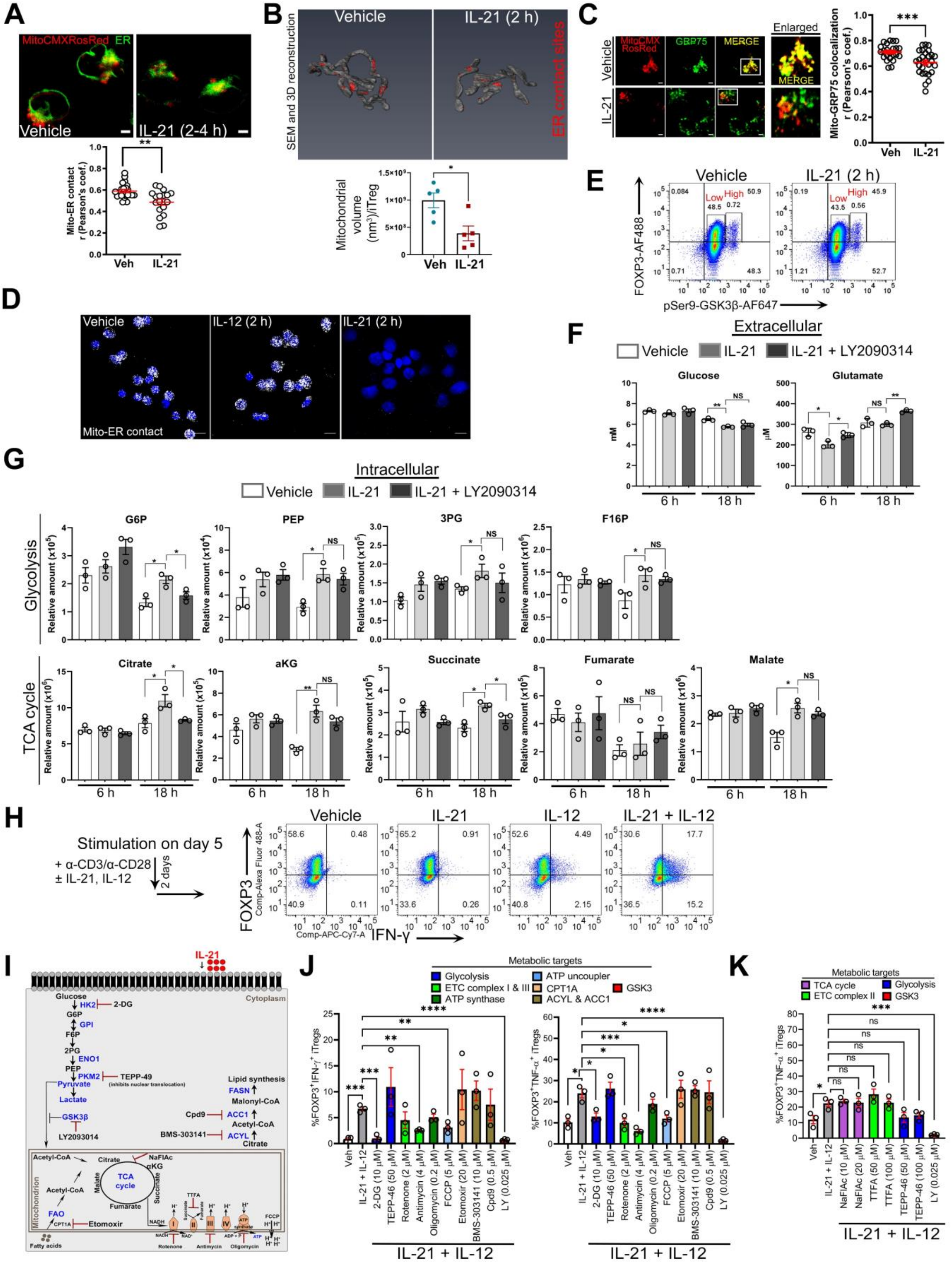
A
MitoCMXRosRed ER
Vehicle
IL-21 (2-4 h)
Mito-ER contact r (Pearson's coef.)
Veh
IL-21
B
Vehicle
IL-21 (2 h)
SEM and 3D reconstruction
ER contact sites
Mitochondrial volume (nm3)/iTreg
Veh
IL-21
C
MitoCMX RosRed
GRP75
MERGE
Enlarged
Vehicle
IL-21
Mito-GRP75 colocalization r (Pearson's coef.)
Veh
IL-21
D
Vehicle
IL-12 (2 h)
IL-21 (2 h)
Mito-ER contact
E
Vehicle
IL-21 (2 h)
FOXP3-AF488
pSer9-GSK3β-AF647
Low
High
F
Extracellular
Vehicle
IL-21
IL-21 + LY2090314
Glucose
Glutamate
6 h
18 h
G
Intracellular
Vehicle
IL-21
IL-21 + LY2090314
Glycolysis
G6P
PEP
3PG
F16P
TCA cycle
Citrate
aKG
Succinate
Fumarate
Malate
Relative amount
6 h
18 h
H
Stimulation on day 5
+ α-CD3/α-CD28
± IL-21, IL-12
2 days
Vehicle
IL-21
IL-12
IL-21 + IL-12
FOXP3
IFN-γ
I
IL-21
Cytoplasm
Glucose
HK2
2-DG
G6P
GPI
F6P
2PG
ENO1
PEP
PKM2
TEPP-49
(inhibits nuclear translocation)
Pyruvate
Lactate
GSK3β
LY2093014
Lipid synthesis
FASN
Malonyl-CoA
Cpd9
ACC1
Acetyl-CoA
BMS-303141
ACYL
Citrate
Acetyl-CoA
NaFIAc
αKG
TCA cycle
Malate
Fumarate
Succinate
TTFA
Mitochondrion
FAO
CPT1A
Etomoxir
Fatty acids
Rotenone
Antimycin
Oligomycin
FCCP
ATP synthase
J
Metabolic targets
Glycolysis
ETC complex I & III
ATP synthase
ATP uncoupler
CPT1A
ACYL & ACC1
GSK3
%FOXP3+IFN-γ+ iTregs
%FOXP3+TNF-α+ iTregs
Veh
IL-21 + IL-12
2-DG (10 μM)
TEPP-46 (50 μM)
Rotenone (2 μM)
Antimycin (4 μM)
Oligomycin (0.2 μM)
FCCP (5 μM)
Etomoxir (20 μM)
BMS-303141 (10 μM)
Cpd9 (0.5 μM)
LY (0.025 μM)
IL-21 + IL-12
K
Metabolic targets
TCA cycle
ETC complex II
Glycolysis
GSK3
%FOXP3+TNF-α+ iTregs
Veh
IL-21 + IL-12
NaFIAc (10 μM)
NaFIAc (20 μM)
TTFA (50 μM)
TTFA (100 μM)
TEPP-46 (50 μM)
TEPP-46 (100 μM)
LY (0.025 μM)
IL-21 + IL-12

**Supplementary Figure 4. IL-21 Dissociates Mitochondria from ER, Resulting in Pyruvate Imbalance and Sensitization to IL-12-induced Inflammatory Response.**

(A) Representative confocal images of live human iTregs loaded with organelle tracker dyes ± IL-21 (2 h) show mito-ER co-localization (yellow) (n = 3). Graph shows mito-ER co-localization in iTregs (n = 22 vs. n = 18) as measured by Pearson's correlation coefficient (Pearson's coef.); scale bar, 5 µm.

(B) Representative 3D reconstructed SEM images of iTreg show ER contact with mitochondria (red) (n = 5). Graph shows mitochondrial volume of per mitochondrion (n = 5 per group).

(C) Representative confocal images of iTregs show GRP75 co-localization with mitochondria (yellow). Graph shows mito-GRP75 co-localization (n = 23 vs. n = 28) as calculated by Pearson's correlation coefficient (Pearson's coef.); scale bar, 2 µm.

(D) Representative PLA images of iTregs show mito-ER contact ± IL-12 or IL-21; scale bar, 5 µm (n = 3).

(E) Representative dot plots show percentage of $FOXP3^+$ pSer-$GSK3\beta^+$ iTregs ± IL-21 (n = 4).

(F and G) Extra- and intra-cellular metabolites from iTregs cultured in normal conditions ± IL-21 (100 ng/ml), LY2090314 GSK3 inhibitor (0.025 µM) or both (n = 3).

(H) Representative dot plots show percentage of $FOXP3^+$ IFN-$\gamma^+$ iTregs ± IL-21 and IL-12 (n = 3).

(I) Schematic illustrates IL-21-induced metabolic processes and established inhibitors of these metabolic pathways: 2-DG, TEPP-46 (inhibits nuclear translocation of PKM2), rotenone (ETC complex I inhibitor), Thenoyltrifluoroacetone (TTFA, ETC complex II inhibitor), antimycin A (ETC complex III inhibitor), oligomycin (ETC complex V or ATP synthase inhibitor), FCCP (ATP uncoupler), etomoxir (inhibitor of CPT1A-mediated FAO), BMS-303141 (acetyl-CoA lyase [ACYL] inhibitor), Cpd9 (acetyl-CoA decarboxylase 1 [ACC1] inhibitor), LY2090314,and NaFlAc (sodium fluoroacetate) (inhibitor of TCA cycle enzymes).

(J and K) Percentage of IL-21 and IL-12-stimulated iTregs that are $FOXP3^+$ IFN-$\gamma^+$ or $FOXP3^+$ TNF-$\alpha^+$ after exposure to indicated inhibitors.

Data represents mean ± SEM. * $p < 0.05$, ** $p < 0.01$, *** $p < 0.001$, and NS (not significant), using two-tailed Student's t test or one-way ANOVA followed by Bonferroni test for multiple comparisons.

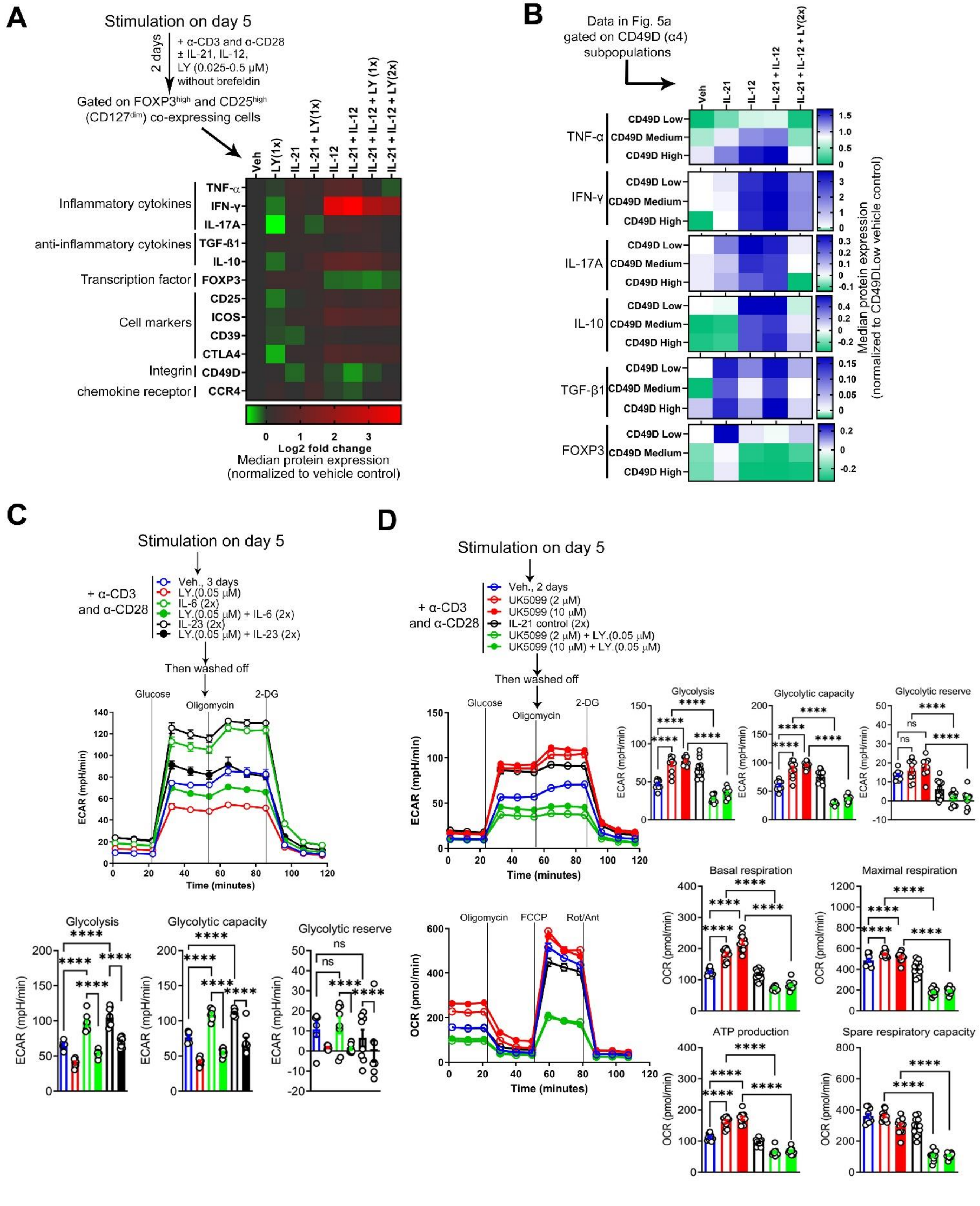
A
Stimulation on day 5
2 days
+ α-CD3 and α-CD28
± IL-21, IL-12,
LY (0.025-0.5 μM)
without brefeldin
Gated on FOXP3high and CD25high
(CD127dim) co-expressing cells
Veh
LY(1x)
IL-21
IL-21 + LY(1x)
IL-12
IL-21 + IL-12
IL-21 + IL-12 + LY (1x)
IL-21 + IL-12 + LY(2x)
Inflammatory cytokines
TNF-α
IFN-γ
IL-17A
anti-inflammatory cytokines
TGF-ß1
IL-10
Transcription factor
FOXP3
Cell markers
CD25
ICOS
CD39
CTLA4
Integrin
CD49D
chemokine receptor
CCR4
0 1 2 3
Log2 fold change
Median protein expression
(normalized to vehicle control)
B
Data in Fig. 5a
gated on CD49D (α4)
subpopulations
Veh
IL-21
IL-12
IL-21 + IL-12
IL-21 + IL-12 + LY(2x)
TNF-α
IFN-γ
IL-17A
IL-10
TGF-β1
FOXP3
CD49D Low
CD49D Medium
CD49D High
Median protein expression
(normalized to CD49DLow vehicle control)
C
Stimulation on day 5
+ α-CD3
and α-CD28
Veh., 3 days
LY.(0.05 μM)
IL-6 (2x)
LY.(0.05 μM) + IL-6 (2x)
IL-23 (2x)
LY.(0.05 μM) + IL-23 (2x)
Then washed off
Glucose
Oligomycin
2-DG
ECAR (mpH/min)
Time (minutes)
Glycolysis
Glycolytic capacity
Glycolytic reserve
ECAR (mpH/min)
****
ns
D
Stimulation on day 5
+ α-CD3
and α-CD28
Veh., 2 days
UK5099 (2 μM)
UK5099 (10 μM)
IL-21 control (2x)
UK5099 (2 μM) + LY.(0.05 μM)
UK5099 (10 μM) + LY.(0.05 μM)
Then washed off
Glucose
Oligomycin
2-DG
ECAR (mpH/min)
Time (minutes)
Glycolysis
Glycolytic capacity
Glycolytic reserve
Oligomycin
FCCP
Rot/Ant
OCR (pmol/min)
Time (minutes)
Basal respiration
Maximal respiration
ATP production
Spare respiratory capacity
OCR (pmol/min)

**Supplementary Figure 5. Methyl Pyruvate Supplementation Mirrors LY2090314 Treatment and Suppresses the Metabolic Basis of Inflammatory iTregs.**

(A and B) Representative heatmaps show $log_2$ fold change of normalized median protein expression, as determined by CYTOF analysis of $FOXP3^{high}$ $CD25^{high}$ IL-7 receptor α-chain $(CD127)^{negative}$ iTregs (A) and analysis of $CD49D^{high,\ medium\ or\ low}$ iTregs (B) (n = 3).

(C) Representative ECAR profile of iTregs ± IL-6 or IL-23 (200 ng/ml), LY2090314 (n = 3). Bar graphs show calculated glycolysis, glycolytic capacity, and glycolytic reserve; mean ± SEM from 10-12 technical replicates.

(D) Representative ECAR (top) and OCR (bottom) profiles of iTregs ± UK5099, IL-21 (200 ng/ml) or UK5099 ± LY2090314 (n = 3). Bar graphs show calculated glycolysis, glycolytic capacity, and glycolytic reserve (top), and basal respiration, ATP production, maximal respiration, and spare respiratory capacity (bottom); mean ± SEM from 10-12 technical replicates.

Data represents mean ± SEM. **** $p < 0.0001$ and ns (not significant), using one-way ANOVA followed by Tukey or Bonferroni test for multiple comparisons.

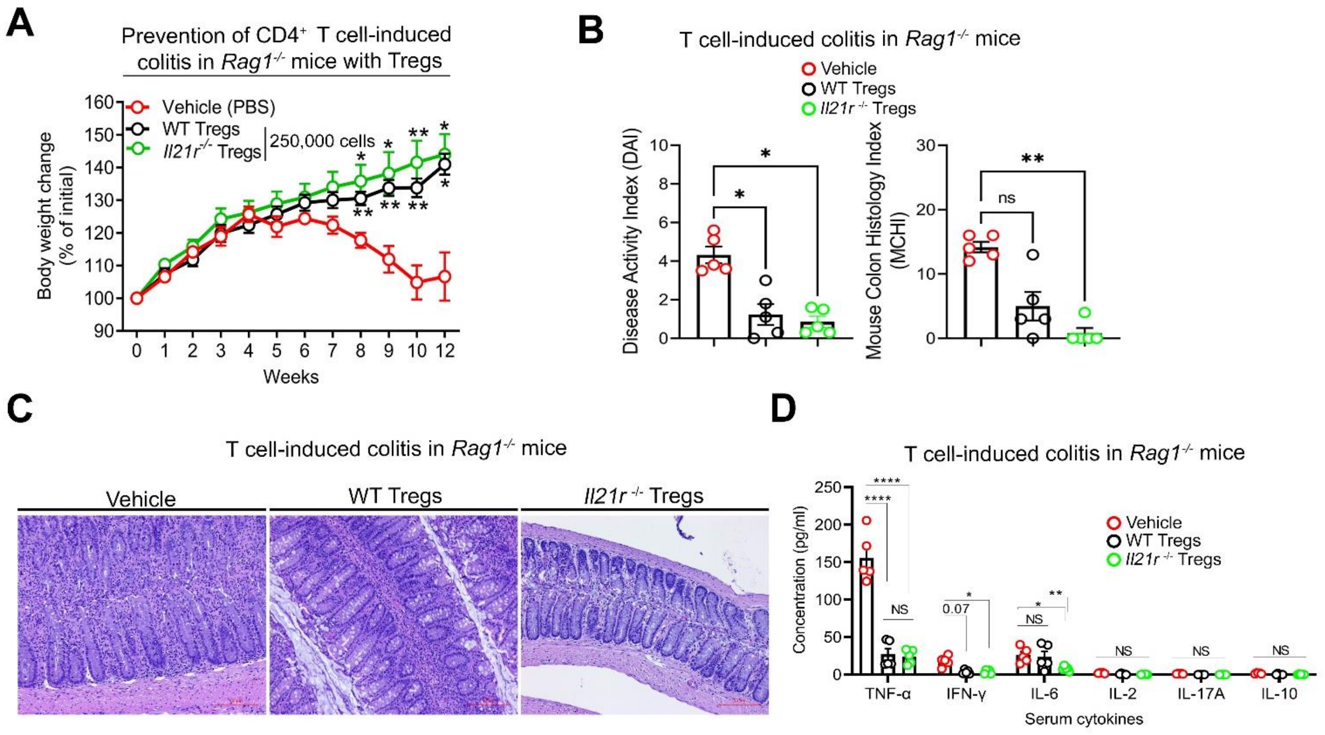


**Supplementary Figure 6. IL-21-induced Metabolic Genes are Enriched in Refractory Human IBD, and IL-21R-deficient Tregs Effectively Lessen CD4$^+$ T Cell-induced Colitis in Mice.**

(A) Change in body weight of mice between week 1 to week 12 during naïve CD4$^+$ T cell-induced colitis progression in *Rag1*$^{-/-}$ mice (n = 5 mice per group).

(B) DAI (top), and MCHI (bottom) of treated colitis mice (n = 5 mice per group). MCHI was assessed by a blinded pathologist.

(C) H&E staining of colon sections of treated colitis mice after 12 weeks; scale bar, 50 µm.

(D) Serum cytokine expression analysis of treated colitis mice on week 12.

Data represents mean ± SEM. * $p < 0.05$, ** $p < 0.01$, and **** $p < 0.0001$, using two-way ANOVA followed by Bonferroni test (A), non-parametric Kruskal-Wallis test followed by Dunn's multiple comparisons (B), and multiple unpaired t tests (D).
